# OpenFold: Retraining AlphaFold2 yields new insights into its learning mechanisms and capacity for generalization

**DOI:** 10.1101/2022.11.20.517210

**Authors:** Gustaf Ahdritz, Nazim Bouatta, Christina Floristean, Sachin Kadyan, Qinghui Xia, William Gerecke, Timothy J O’Donnell, Daniel Berenberg, Ian Fisk, Niccolò Zanichelli, Bo Zhang, Arkadiusz Nowaczynski, Bei Wang, Marta M Stepniewska-Dziubinska, Shang Zhang, Adegoke Ojewole, Murat Efe Guney, Stella Biderman, Andrew M Watkins, Stephen Ra, Pablo Ribalta Lorenzo, Lucas Nivon, Brian Weitzner, Yih-En Andrew Ban, Peter K Sorger, Emad Mostaque, Zhao Zhang, Richard Bonneau, Mohammed AlQuraishi

## Abstract

AlphaFold2 revolutionized structural biology with the ability to predict protein structures with exceptionally high accuracy. Its implementation, however, lacks the code and data required to train new models. These are necessary to (i) tackle new tasks, like protein-ligand complex structure prediction, (ii) investigate the process by which the model learns, which remains poorly understood, and (iii) assess the model’s generalization capacity to unseen regions of fold space. Here we report OpenFold, a fast, memory-efficient, and trainable implementation of AlphaFold2. We train OpenFold from scratch, fully matching the accuracy of AlphaFold2. Having established parity, we assess OpenFold’s capacity to generalize across fold space by retraining it using carefully designed datasets. We find that OpenFold is remarkably robust at generalizing despite extreme reductions in training set size and diversity, including near-complete elisions of classes of secondary structure elements. By analyzing intermediate structures produced by OpenFold during training, we also gain surprising insights into the manner in which the model learns to fold proteins, discovering that spatial dimensions are learned sequentially. Taken together, our studies demonstrate the power and utility of OpenFold, which we believe will prove to be a crucial new resource for the protein modeling community.

## 1 Introduction

Predicting protein structure from sequence has been a defining challenge of biology for decades ([1], [2]). Building on a line of work applying deep learning to co-evolutionary information encoded in multiple sequence alignments (MSAs) ([3], [4], [5], [6], [7], [8]) and homologous structures ([9], [10]), AlphaFold2 ([11]) has arguably solved the problem for natural proteins with sufficiently deep MSAs. The model has been made available to the public with DeepMind’s official open-source implementation, which has enabled researchers to optimize AlphaFold2’s prediction procedure and user experience ([12]) and to employ it as a module within novel algorithms, including ones for protein complex prediction ([13]), peptide-protein interactions ([14]), structure ranking ([15]), and more (*e.g.*, [16], [17], [18]). It has also been used to predict the structures of hundreds of millions of proteins ([19], [20], [21])

In spite of its outstanding utility, the official AlphaFold2 implementation omits code for the model’s complex training procedure as well as the computationally expensive data required to run it. This makes it difficult to i) investigate AlphaFold2’s learning behavior and sensitivity to changes in data composition and model architecture and ii) create variants of the model to tackle new tasks. Given the success of AlphaFold2, its many novel components are likely to prove useful for tasks beyond protein structure prediction. For instance, retraining AlphaFold2 using a dataset of protein-protein complexes resulted in AlphaFold2-Multimer ([22]), the state of the art model for predicting structures of protein complexes. Until recently, however, this capability has been exclusive to DeepMind.

To address this shortcoming, we developed OpenFold, a *trainable* open-source implementation of AlphaFold2. We trained OpenFold from scratch using OpenProteinSet [23], our open-source reproduction of the AlphaFold2 training set, matching AlphaFold2 in prediction quality. Apart from new training code and data, OpenFold has several advantages over AlphaFold2: (i) it runs up to three times faster on most proteins, (ii) it uses less memory, allowing prediction of extremely long proteins and multi-protein complexes on a single GPU, and (iii) it is implemented in PyTorch ([24]), the most widely used machine learning framework (AlphaFold2 uses Google’s JAX ([25])). As such, OpenFold can be readily used by the widest community of developers and interfaces with a rich ecosystem of existing machine learning software ([26], [27], [28], [29]).

Taking advantage of our discovery that ∼90% of model accuracy can be achieved in ∼3% of training time, we retrained OpenFold multiple times on specially elided versions of the training set to quantify its ability to generalize to unseen protein folds. Surprisingly, we found the model highly robust even to large elisions of fold space, but its capacity to generalize varied based on the spatial extent of protein fragments and folds. We observed even stronger performance when training the model on more diverse but smaller datasets, some as small as 1,000 experimental structures. Next, we used OpenFold to understand how the model learns to fold proteins, focusing on the geometric characteristics of predicted structures during intermediate stages of training, and identified multiple distinct phases of behavior. Specifically, by analyzing predicted structures at multiple resolutions and decomposing them into secondary and tertiary elements, we found that OpenFold learns spatial dimensions, secondary structure elements, and tertiary scales in a staggered manner. Taken together, these results yield fundamental new insights into the learning behavior of AlphaFold2-type models and provide new conceptual and practical tools for the development of biomolecular modeling algorithms.

## 2 Results

### Newly trained OpenFold matches AlphaFold2 in accuracy

OpenFold reproduces the AlphaFold2 model architecture in full, without any modifications that could alter its internal mathematical computations. This results in perfect interoperability between OpenFold and AlphaFold2, enabling use of the original AlphaFold2 model parameters within OpenFold and vice versa. To verify that our OpenFold implementation recapitulates all aspects of AlphaFold2 training, we used it to train a new model from scratch. OpenFold/AlphaFold2 training requires a collection of protein sequences, MSAs, and structures. As the AlphaFold2 MSA database has not been publicly released, we used OpenProteinSet [23], a replication of the AlphaFold2 training dataset that substitutes newer versions of sequence databases where available. Starting from approximately 15 million Uni-Clust30 ([30]) MSAs, we selected approximately 270,000 diverse and deep MSAs to form a “self-distillation” set; such sets are used to augment experimental training data with high-quality predictions. We predicted protein structures for all MSAs in this set using AlphaFold2 and combined them with approximately 132,000 unique (640,000 non-unique) experimental structures from the Protein Databank ([31]) to form the OpenFold training data set. During training on self-distillation proteins, residues with a low AlphaFold2 confidence score (< 0.5 pLDDT) were masked. Our validation set consisted of nearly 200 structures from CAMEO ([32]), an online repository for continuous quality assessment of protein structure prediction models, drawn over a three-month period ending on January 16, 2022.

From our main training run, we selected seven snapshots to form a collection of distinct (but related) models. During prediction time, these models can generate alternate structural hypotheses for the same protein. To further increase the diversity of this collection, we fine-tuned a second set of models that we branched off from the main model. In this second branch, we disabled the model’s template pipeline, similar to the procedure used for AlphaFold2. Selected snapshots from this branch were added to the pool of final models, resulting in a total of 10 distinct models. Full training details are provided in Appendix C. We summarize the main results of our training experiment in Figure 1. Predictions made by OpenFold and AlphaFold2 on the CAMEO validation set are assessed using the lDDT-C ([33]) metric (Figure 1A) and show very high concordance between OpenFold and AlphaFold2, demonstrating that OpenFold successfully reproduces AlphaFold2. While OpenFold was still training for much of the recent CASP15 competition, our retrospective evaluation shows that the final model achieves parity on CASP15 domains as well (Figure 1). Figure 1C provides a visual illustration of this concordance. Tracking prediction accuracy as a function of training stage (Figure 1D) reveals the remarkable fact that OpenFold achieves ∼90% of its final accuracy in just 1,500 GPU hours (∼3% of training time) and ∼95% in 2,500 GPU hours; total training time is approximately 50,000 GPU hours. This rapid rise in accuracy suggests that training of new OpenFold variants can be accomplished with far less compute than is necessary for full model training, facilitating rapid exploration of model architectures. We take advantage of this fact in our data elision experiments.

**Figure 1:**
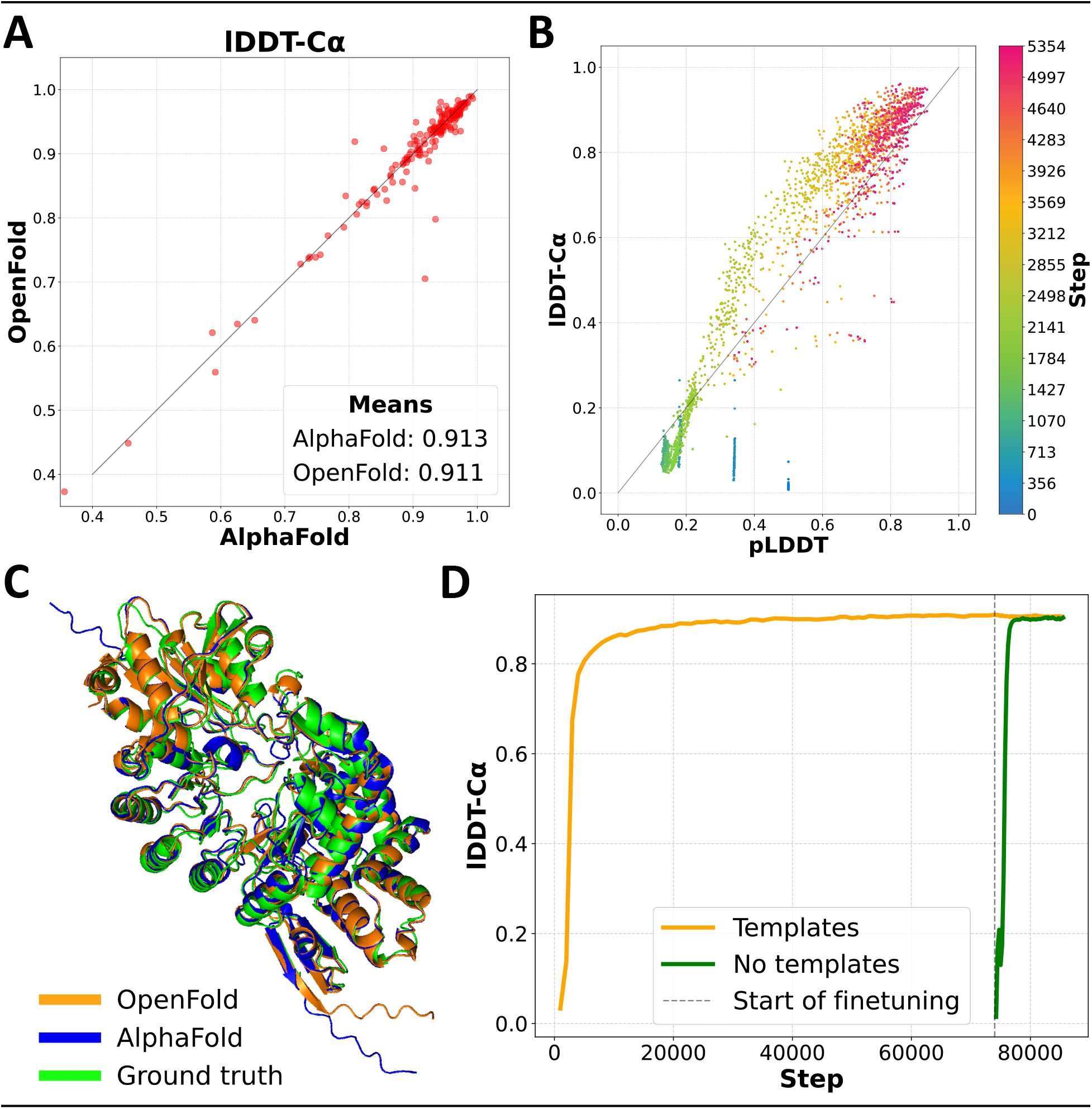
OpenFold matches the accuracy of AlphaFold2. **(A)** Scatter plot of lDDT-C values of AlphaFold and OpenFold predictions on the CAMEO validation set. **(B)** Average pLDDT vs lDDT-C of OpenFold predictions on the CAMEO set during the early stage of training. OpenFold is initially overconfident but quickly becomes underconfident, gradually converging to accurate confidence estimation. **(C)** Predictions by OpenFold and AlphaFold2 overlayed with an experimental structure of *S. tokunonesis* TokK protein ([34]; PDB accession code: 7KDX_A). **(D)** Average lDDT-C for OpenFold computed over the training set during the course of training. The template-free branch is shown in green, the template-utilizing one in orange, and the initial-training/fine-tuning boundary in black. Template-free accuracy is initially poor because the exponential moving average of the weights used for validation was being reinitialized.

AlphaFold2 training is broadly split into two phases, an initial training phase and a more computationally intensive fine-tuning phase. In the latter, the size of protein fragments used for training is increased to 384 residues and an additional loss function that penalizes structural violations (*e.g.* steric clashes) is enabled. By comparing predicted structures between the initial and fine-tuning phases, we find that the second phase has only a modest effect on overall structural quality metrics, even when considering only long proteins greater than 500 residues in length (see Appendix F.1). Instead, the primary utility of fine-tuning appears to be to resolve violations of known chemical constraints. In our training experiments, this occurs quickly after the beginning of fine-tuning, suggesting that elided fine-tuning runs can be used with minimal impact on prediction quality.

In addition to prediction accuracy, we also tracked pLDDT as a function of training stage. pLDDT is the model’s estimate of the lDDT-C of predicted structures and serves as its primary confidence metric. We find that pLDDT is well correlated with true lDDT early in training, albeit initially over-confident in its self-assessment and later entering a phase of under-confidence (Figure 1B). It is notable that the model is capable of assessing the quality of its own predictions early on in training, when its overall predictive capacity remains very limited.

### OpenFold can achieve high accuracy using tiny training sets

Having established the equivalency of AlphaFold2 and OpenFold, we set out to understand properties of the architecture, starting with its data efficiency. AlphaFold2 was trained using ∼132,000 protein structures from the PDB, the result of decades of painstaking and expensive experimental structure determination efforts. For other molecular systems for which AlphaFold2-style models may be developed, data is far more sparse; *e.g.,* the PDB contains only 1,664 RNA structures. We wondered whether the high accuracy achieved by AlphaFold2 in fact depended on its comparatively large training set, or if it is possible to achieve comparable performance using less data. Were the latter to be true, it would suggest broad applicability of the AlphaFold2 paradigm to molecular problems. To investigate this possibility, we performed a series of OpenFold training runs in which we used progressively less training data, assessing model accuracy as a function of training set size.

Our first set of tests randomly subsample the original training data to 17,000, 10,000, 5,000, 2,500, 2,000, and 1,000 protein chains. We used each subsampled set to train OpenFold for at least 7,000 steps, through the initial rapid rise phase to early convergence. To avoid information leakage from the full training set, we did not use self-distillation, putting the newly trained models at a disadvantage relative to the original OpenFold. We trained models with and without structural templates. In all other regards, training was identical to that of the standard OpenFold model. Model accuracy (assessed using lDDT-C) is plotted as a function of training step in Figure 2A, with colors indicating size of training set used.

**Figure 2:**
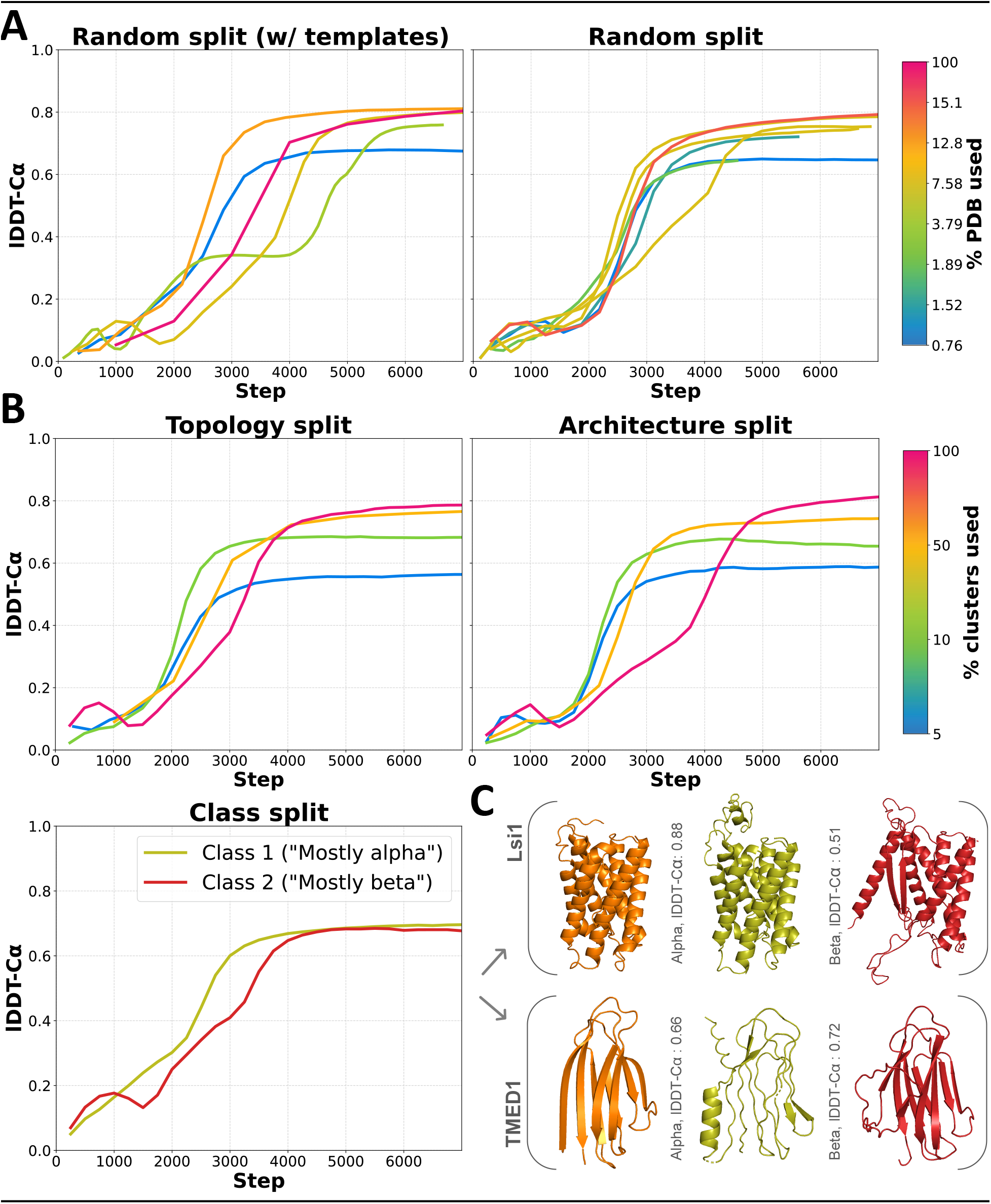
OpenFold generalization capacity on elided training sets. **(A)** Validation set lDDT-C as a function of training step for models trained on elided training sets (10k random split repeated 3x). **(B)** Same as (A) but for CATH-stratified dataset elisions. Validation sets vary across stratifications and are not directly comparable. **(C)** Experimental structures (orange) and mainly alpha-trained (yellow) and mainly beta-trained (red) predictions of largely helical Lsi1 (top) and beta sheet-heavy TMED1 (bottom).

We find that merely protein chains—about of all training data (yellow curves)—suffice to reach essentially the same initial lDDT-C as a model trained on the full training set (pink curve). After 20,000 steps (not pictured), the full data model reaches a peak lDDT-C of 0.83, while after 7,000 steps, the 10,000-sample model has already exceeded 0.81 lDDT-C . Although performance gradually degrades as training set size decreases further, we find that all models are surprisingly performant, even ones trained on our smallest subsample of 1,000 protein chains, corresponding to just 0.76% of the full training set. In fact, this model reaches an lDDT-C of 0.64, exceeding the median lDDT-C of 0.62 achieved at CASP13 by the first AlphaFold, the best performing model at the time.

Comparing the accuracies of models trained with and without templates, we find that templates on average contribute little to prediction quality even in the low-data setting. This is consistent with the original AlphaFold2 ablation studies which showed that templates have a minimal effect except when MSAs are shallow or entirely absent.

### OpenFold generalizes to unseen regions of fold space

Randomly subsampling the OpenFold training set, as in the preceding analysis, reduces the quantity of the training data used but not necessarily its overall diversity. In molecular modeling tasks, the data available for training often does not reflect the underlying diversity of the molecular system being modeled, due to biases in the scientific questions pursued, experimental assays available, *etc*. To assess OpenFold’s capacity to generalize to out-of-distribution data, we subsample the training set in a structurally stratified manner such that entire regions of fold space are excluded from training but retained for model assessment. Multiple structural taxonomies for proteins exist, including the hierarchical CATH ([35], [36]) and SCOP ([37]) classification systems. For this task we use CATH, which assigns protein domains—in increasing order of specificity—to a (C)lass, (A)rchitecture, (T)opology, and (H)omologous superfamily. Domains with the same homologous superfamily (H) classification may differ superficially but have highly similar structural cores. Our preceding analysis can be considered to structurally stratify data at the H level. For the present analysis, we stratify data further, holding out entire topologies (T), architectures (A), and classes (C).

We start by filtering out protein domains that have not been classified by CATH, leaving ourselves with ∼440,000 domains spanning 1,385 topologies, 42 architectures, and 4 classes. For the topology stratification, we randomly sample 100 topologies and remove all associated chains from the training set. We construct a validation set from the held out topologies by sampling one representative chain from each. We also construct successively smaller training sets from shrinking fractions of the remaining topologies, including a training set that encompasses all of them. We follow an analogous procedure for architectures except that in this case, the validation set consists of 100 chains randomly selected from 5 architectures (20 per architecture). For class-based stratification, the validation set comprises domains that are neither in the mainly alpha nor mainly beta classes, hence enriching for domains with high proportions of both SSEs. For training, we construct two sets, one corresponding exclusively to the mainly alpha class and another to the mainly beta class—this enables us to ascertain the capability of models trained largely on either alpha helices or beta sheets to generalize to proteins containing both. For all stratifications, we train OpenFold to early convergence (∼7,000 steps) from scratch. To prevent leakage of structural information from held out categories, all runs are performed without templates. We plot model accuracies as a function of training step in Figure 2B, with colors indicating the fraction of categories retained during training for each respective level of the CATH hierarchy.

As expected, removing entire regions of fold space has a more dramatic effect on model performance than merely reducing the size of the training set. For example, retaining of topologies for training (green curve in Figure 2B, topology split), which corresponds to ∼6,400 unique chains, results in a model less performant than one containing 5,000 randomly selected chains (green curve in Figure 2A, no templates). However, even in the most severe elisions of training set diversity, absolute accuracies remain unexpectedly high. For instance, the training set containing of topologies (2,000 chains) still achieves an lDDT-C near 0.6, comparable again to the first AlphaFold, which was trained on over 100,000 protein chains. Similarly, the training set for the smallest architecture-based stratification only contains domains from one architecture (out of 42 that cover essentially the entirety of the PDB), yet it peaks near 0.6 lDDT-C . Most surprisingly, the class-stratified models, in which alpha helices or beta sheets are almost entirely absent from training, achieve very high lDDT-C scores of >0.7 on domains containing both alpha helices and beta sheets. These models likely benefit from the comparatively large number of unique chains in their training sets—15,400 and 21,100 for alpha helix- and beta sheet-exclusive sets, respectively. It should also be noted that the mainly alpha and mainly beta categories do contain small fractions of beta sheets and alpha helices, respectively (see Supplementary Figure 9). Despite these caveats, the model is being tasked with a very difficult out-of-distribution generalization problem in which unfamiliar types of SSEs (from the perspective of the training set) have to essentially be inferred with minimal quantities of corresponding training data. Taken together, these results show that the AlphaFold2 architecture is capable of remarkable feats of generalization.

To better understand the behavior of class-stratified models, we analyzed the structures of two protein domains, one composed almost exclusively of alpha helices (rice Lsi1 aquaporin domain ([38])) and another of beta sheets (human TMED1 domain ([39])), as they are predicted by models trained on the mainly alpha or mainly beta datasets. In the top row of Figure 2C, we show an experimental structure (orange) for Lsi1^1^ along with predictions made by the mainly alpha-trained model (purple) and mainly beta-trained model (green). In the bottom row we show similar figures for TMED1^2^. Predictably, the mainly alpha-trained model accurately predicts the alpha helices of Lsi1 but fails to properly form beta sheets for TMED1 and incorrectly adopts a small alpha helix in part of the structure. The mainly beta-trained model has the opposite problem: its Lsi1 prediction contains poorly aligned helices and an erroneous beta sheet, but TMED1 is reasonably well predicted. Notably, however, neither fails catastrophically. Regions corresponding to the beta sheets of Lsi1 are predicted by the mainly alpha model with approximately the right shape, except that their atomic coordinates are not sufficiently precise to enable DSSP to classify them as beta sheets.

### Generalization capacity is scale-dependent

OpenFold’s surprising capacity for generalization across held out regions of fold space suggests that it is somewhat indifferent to the diversity of the training set at the global fold level. Instead, the model appears to learn how to predict protein structures from local patterns of MSA/sequence-structure correlations—fragments, secondary structure elements, individual residues, and so on—rather than from global fold patterns captured by CATH. This raises the possibility that the model’s capacity for generalization depends on the spatial scale of the prediction task. To directly test this hypothesis, we assessed model accuracy on protein fragments of increasing length using both the GDT_TS and lDDT-Cα metrics as a function of the fraction of the training set retained for topology- and architecture-stratified models (Figure 3). Note that lDDT-Cα is less sensitive to global fit than GDT_TS. To make results directly comparable between different stratifications, we used a common validation set derived from CAMEO. This validation set likely contains domains from all CATH categories and may thus overestimate accuracies.

**Figure 3:**
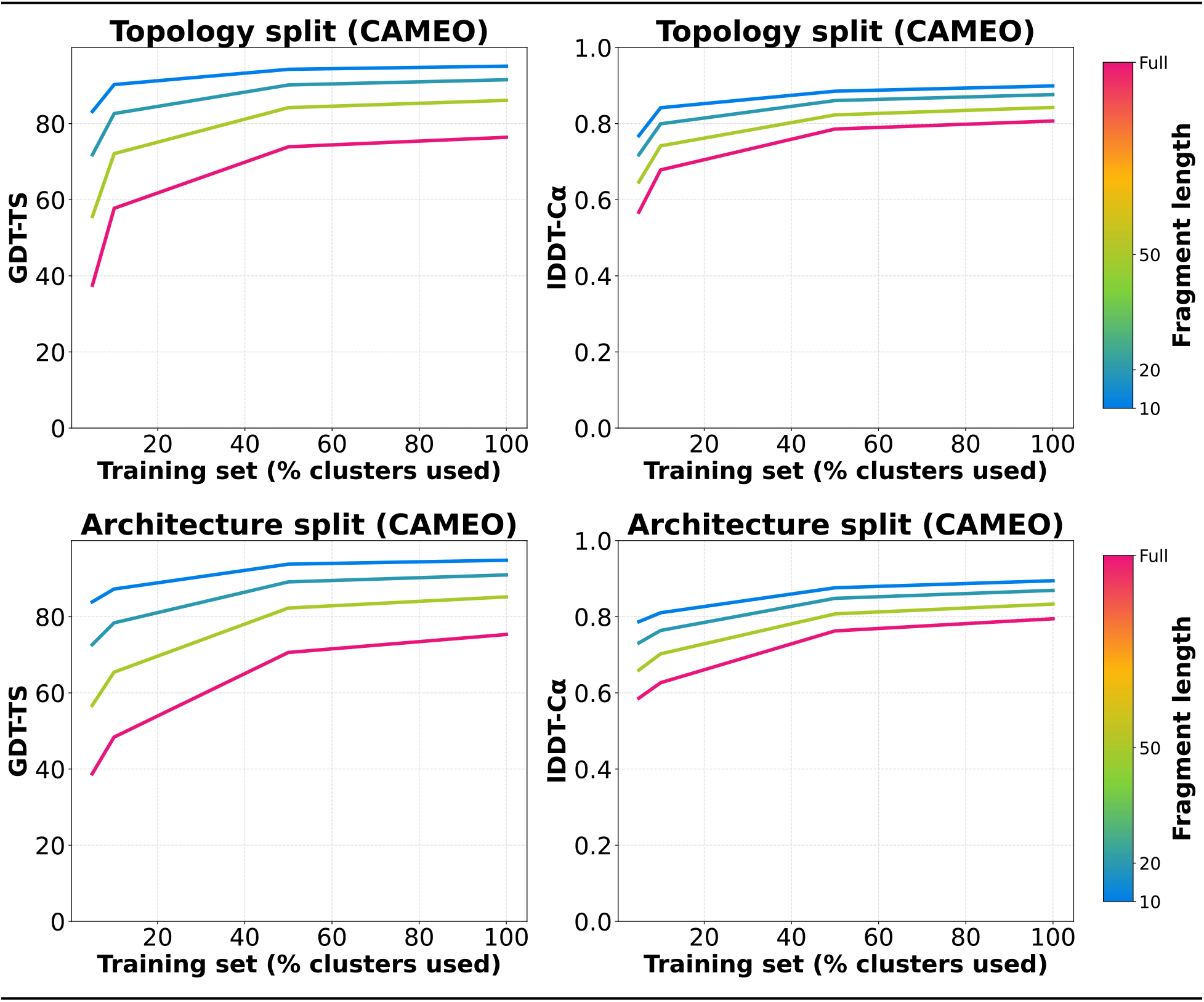
Reduced dataset diversity disproportionately affects global structure. Mean GDT-TS and lDDT-C of non-overlapping protein fragments from CAMEO validation set as a function of the percentage of CATH clusters in elided training sets. Data for both topology and architecture elisions are included. Fragmenting procedure is the same as that described in Figure 6A.

We observe that fragments of all lengths are better predicted when more data is used for training. However, the relative gains seen by larger fragments and whole domains (pink) far exceed those seen by smaller fragments (blue), consistent with the hypothesis that generalization capacity is length-dependent. In particular, it indicates that local structure can in fact be robustly learned from highly elided data sets, while global structure is more dependent on broad representation of fold diversity in the training set.

### OpenFold learns a sequence of approximate PCA projections

We have shown that AlphaFold2 very readily achieves high accuracy, even in the absence of diverse training data. Exactly how it does so, however, is poorly understood. In the following sections, we shed light on this fundamental question by analyzing structures predicted by OpenFold as it progresses through training. We focus in particular on the first five thousand steps of training, during which the model experiences rapid gains in accuracy. Overall, we find that AlphaFold2 learns in a highly staggered fashion, perfecting individial spatial dimensions of predicted structures before others, simple secondary structure elements before rarer ones, and local structure before global structure.

First, we find a remarkably consistent progression in the dimensionality of predicted structures, best visualized in animations we provide in the supplement^3^. At first, predictions are point-like and zero-dimensional, as the model is initialized to place all residues at the origin. Early on, structures extend along a single axis, remaining approximately one-dimensional. A few hundred training steps later, these tubular predictions begin to stretch in an orthogonal dimension to resemble curved, two-dimensional surfaces. In this stage, flattened secondary structure elements are often clearly visible. Once the two-dimensional extent of the final structure is nearly fully realized, predicted structures begin to inflate and acquire volume along a third orthogonal dimension, arriving at reasonably accurate backbone structures. Finally, after global geometry stabilizes, secondary structure elements begin to acquire precise shape and atomic detail. Subsequent epochs make largely minor, local revisions to secondary structure elements, which we analyze in a later section. We note that this progression differs markedly from that obtained when analyzing structures predicted from intermediate layers of a fully trained model, which are all generally globular and three-dimensional in nature with at least partially-formed secondary structure elements ([11]).

During our analysis of structures predicted at intermediate stages of training, we observed that predictions in the 2D phase contain patterns reminiscent of two-dimensional projections of the secondary structure elements of the final globular structure. This fact contributes to a distinct impression that predictions don’t simply undergo phases of dimensionality, but that during each phase they “grow” predominantly along the new dimension corresponding to that phase. More precisely, the model appears to learn to generate successive lower dimensional PCA projections of the true 3D structures, first learning a 1D PCA projection of the final structure, then a 2D PCA projection, and finally the full 3D structure.

To investigate this learning hypothesis, we created 1D and 2D PCA projections of predicted structures at each timestep and compared them to the full 3D prediction at training step 5,000, at the end of the rapid rise in accuracy, using the translationally- and rotationally-invariant distance-based root mean square deviation (dRMSD) metric ([40]). Results are shown in Figure 4A. Before the model exits the 1D and 2D prediction phases, the full (non-projected) predictions (“3D”) and their lower dimensional projections (“1D” and “2D”) are almost indistinguishable from each other, as expected; at these stages, as we previously described, predicted structures are essentially one- or two-dimensional. Thereafter, as the predictions gain dimensions, their dRMSDs to the final structure diverge from those of the lower dimensional projections. Remarkably, much of the overall drop in dRMSD occurs before the 2D and 3D projections diverge, indicating that the model is improving its accuracy in lower dimensions before moving on to higher dimensions. It does not perfect each projection before transitioning, however. Figure 4B shows a similar experiment to 4A except low-dimensional projections for each training iteration are compared to corresponding low-dimensional projections of the final prediction at step 5,000. In the extreme, if the model were learning perfect low-dimensional PCA projections of the final 3D structure at the end of each low-dimensional training phase, the 2D projections of the intermediate and final predictions would match exactly at the end of the 2D phase. Additionally, since the 3D prediction is essentially flat at the end of the 2D phase, its dRMSD with respect to the 3D structure should be high. Such a separation is not visible in Figure 4B; instead, all three projections converge to their final counterparts at nearly the same time. This suggests that while the model learns crude approximations of low-dimensional PCA projections during each phase, its learning is largely continuous, with all spatial dimensions continuing to be refined until the end. However, the degree to which the dominant dimensions continue to be refined diminishes over time relative to less dominant ones.

**Figure 4:**
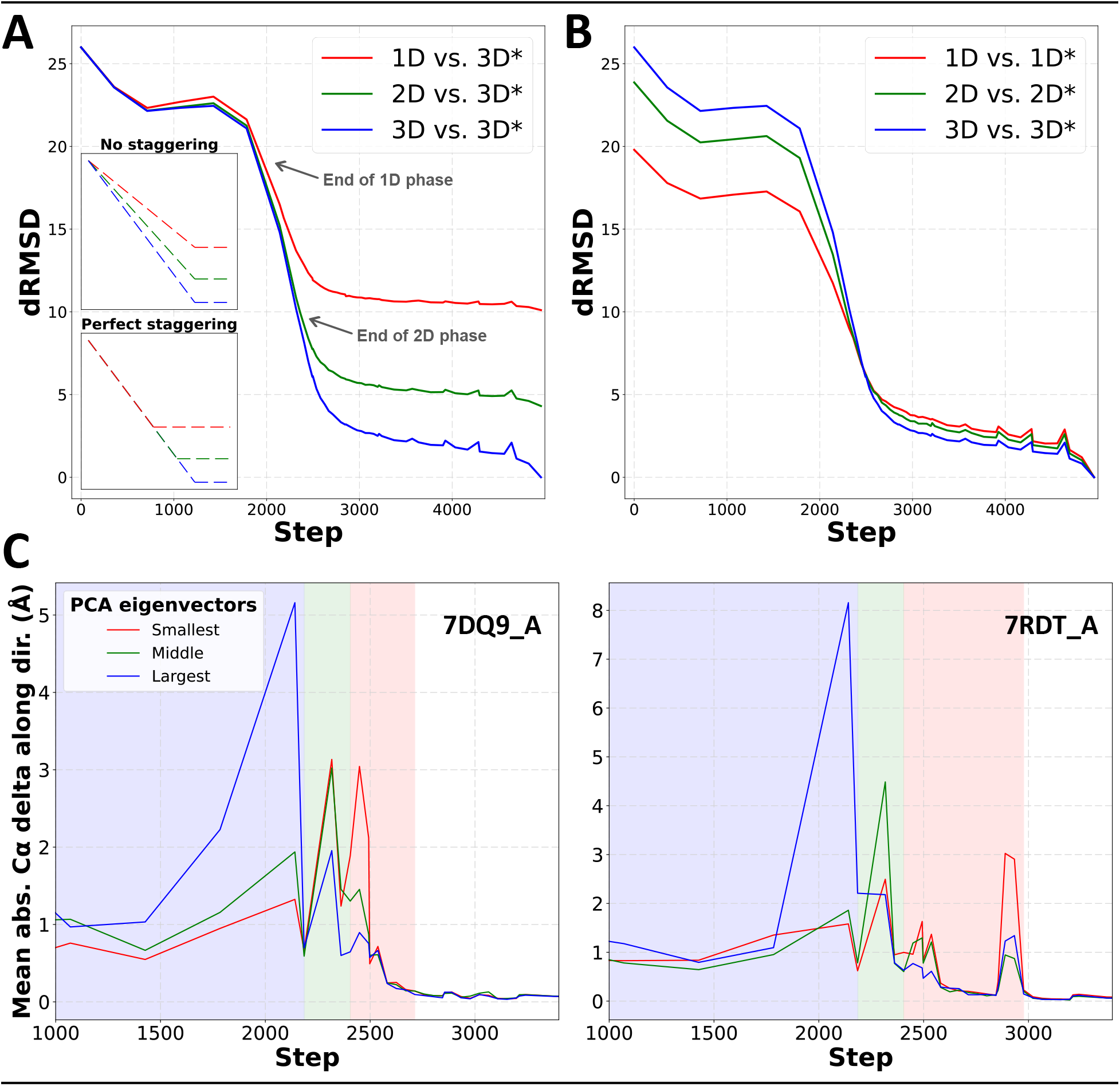
Early predictions crudely approximate lower-dimensional PCA projections. **(A)** Mean dRMSD, as a function of training step, between low-dimensional PCA projections of predicted structures and the final 3D prediction at step 5,000 (denoted by *). Averages are computed over the CAMEO validation set. Insets show idealized behavior corresponding to unstaggered, simultaneous growth in all dimensions and perfectly staggered growth. Empirical training behavior more closely resembles the staggered scenario. **(B)** Low-dimensional projections as in (A) compared to projections of the final predicted structures at step 5,000. **(C)** Mean displacement, as a function of training step, of C atoms along the directions of their final structure’s PCA eigenvectors. Results are shown for two individual proteins (PDB accession codes 7DQ9_A and 7RDT_A). Shaded regions correspond loosely to the “1D,” “2D,” and “3D” phases of dimensionality.

To better assess this phenomenon and gain a finer-grained view of the progress that occurs during each phase, we analyzed the movement of atoms along the directions of the final prediction’s principal components as a function of training step. Because predicted structures and their associated principal components are in principle quite mobile over the course of training, this movement is difficult to characterize precisely. Nonetheless, we devised the following scheme to estimate it. Given predicted structures in chronological order, for each in, we align the th structure to the th structure sequentially. Then, for every pair of consecutive predictions, we compute the absolute value of the displacement of each C atom along the directions of the three principal components of the final prediction. In Figure 4C we sequentially plot these displacements for two CAMEO proteins. As before, there is not a perfect separation between phases, and discernible motion occurs in all three directions in every phase. This is compounded by the fact that predictions in the “2D phase” are generally not flat two-dimensional sheets but instead exhibit some degree of curvature and thus produce spurious movement in the third dimension. Nevertheless, both proteins show clearly differentiated spikes in each phase corresponding to rapid expansion in the phases’ respective dimensions. Furthermore, each protein nears its maximum spatial extent in each principal direction during the corresponding phase. After the 1D phase, for example, growth along the direction of the first principal component is greatly subdued. Intuitively, the model appears to exhaust the easiest gains in the most dominant dimensions before proceeding to the less dominant ones, making relatively minor adjustments in previous dimensions thereafter. It is worth noting that, if intuitive, this process is only possible because—unlike previous generations of protein folding models like AlphaFold1 ([7]) or RGN ([41])—AlphaFold2 is not constrained by strict physical priors at this stage, allowing it to output low-dimensional structures with numerous steric clashes that exhibit gross violations of basic chemical laws. This freedom appears to be an important ingredient in AlphaFold2’s success; in the original AlphaFold2 paper, it is observed without further elaboration that enabling a violation loss to penalize steric clashes and non-physical bond lengths in the early stages of training has a strongly destabilizing effect ([11])

### Learning of secondary structure is staggered and multi-scale

The preceding analysis suggests that secondary structure elements (SSEs) are learned subsequent to tertiary structure. We next set out to formally confirm this observation and chronicle the order in which distinct SSEs are learned. For every protein in our validation set and every step of training, we used DSSP ([42]) to identify residues matching the eight recognized SSE states. We treat as ground truth DSSP assignments of residues in the experimental structures, and compute F1 scores as a combined metric of the recall and precision achieved by the model for every type of SSE at various training steps (Figure 5A).

**Figure 5:**
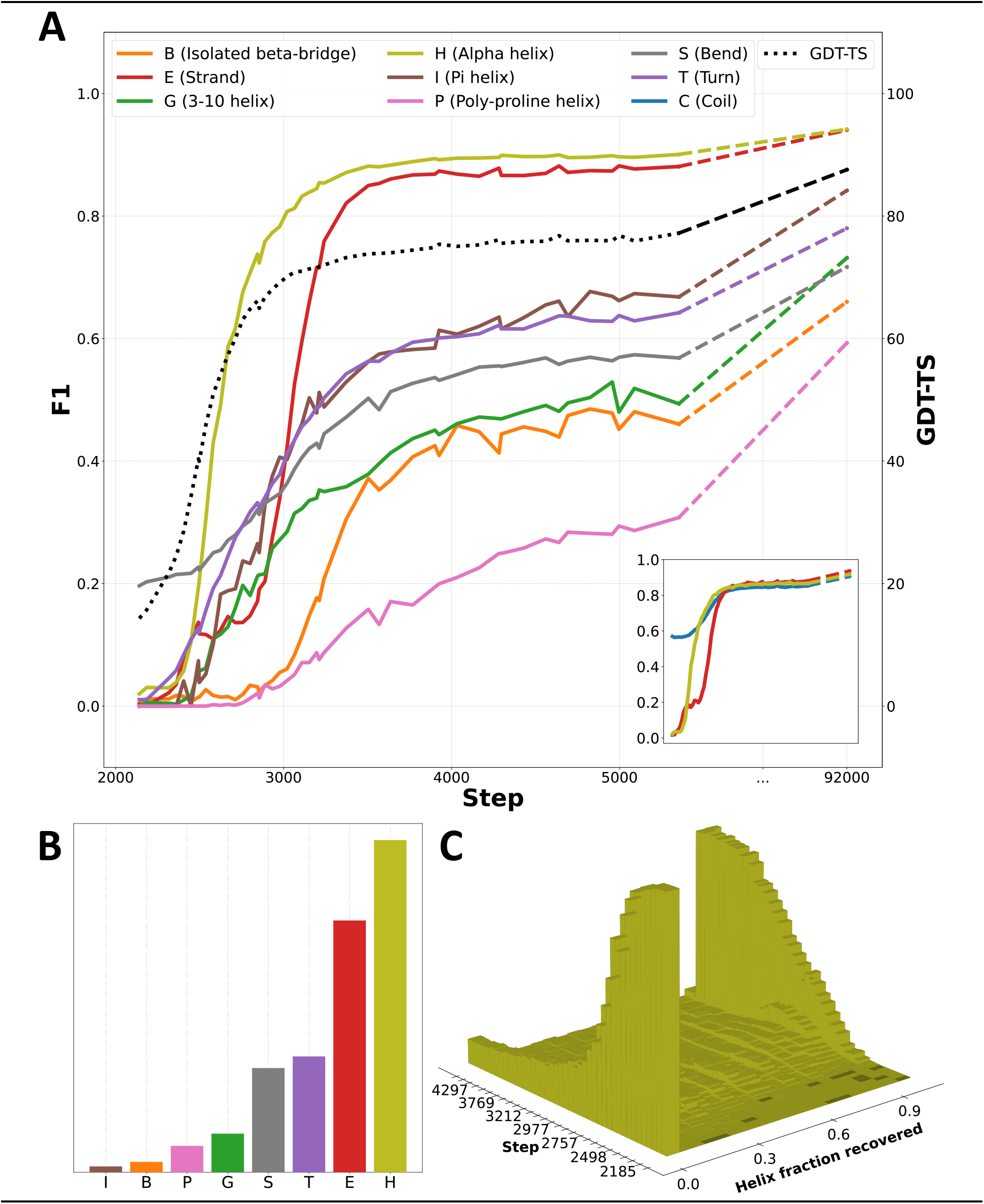
Secondary structure categories are learned in succession. **(A)** F1 scores for secondary structure categories over time. The corner pane depicts the same data using a simplified 3-state assignment (details in Appendix F.3). GDT-TS and final values are also provided. **(B)** Corresponding counts of individual secondary structure assignments. **(C)** Contiguous fractions of individual helices recovered early in training.

We observe a clear sequence in which SSEs are discovered: alpha helices are learned first, followed by beta sheets, followed by less common SSEs. Unsurprisingly, this sequence roughly corresponds to the relative frequencies of SSEs in proteins (Figure 5B), with the exception of uncommon helix variants. As was previously evident, the model’s discovery of SSEs lags that of accurate global structure. For instance, the F1 score for beta sheets (‘E’) only plateaus hundreds of steps after global structural accuracy, as measured by GDT-TS ([43]). This is also clearly visible in our animations of progressive training predictions; for each protein, secondary structure is recognized and rendered properly only after global geometry is essentially finalized.

To investigate the possibility that OpenFold is achieving high alpha helical F1 scores by gradually learning small fragmentary helices, we binned predicted helices by the longest contiguous fraction of the ground-truth helix they recover and plotted the resulting histogram as a function of training step in Figure 5C. Evidently, little probability mass ever accumulates between —helices that are not recovered at all—and 1.0—helices that are completely recovered. This suggests that, at least from the perspective of DSSP, most helices become correctly predicted essentially all at once. This sudden transition coincides with most of the improvement in helix DSSP F1 scores in the early phase of training.

Based on the above observation, we reasoned that as training progresses, OpenFold may first learn to predict smaller structural fragments before larger ones, and that this may be evident on both the tertiary and secondary structure levels. Focusing first on tertiary structure, we assessed the prediction quality, at each training step, of all non-overlapping fragments of length 10, 20, and 50 residues in our validation set. Average GDT-TS values are shown in Figure 6A. Unlike global GDT-TS (pink), which improves minimally in the first 300-400 steps of training, fragment GDT-TS improves markedly during this phase, with shorter fragments showing larger gains. By step 1,000, when the model reaches a temporary plateau, it has learned to predict local structure far better than global structure (GDT-TS > 50 for 10-residue fragments vs GDT-TS < 10 for whole proteins). Soon after, at step 1,800, the accuracy of all fragment lengths including global structure begin to rise rapidly. However, the gains achieved by shorter fragments are smaller than those of longer fragments, such that the gap between 10-residue fragments and whole proteins is much smaller at step 3,000 than 1,800 (GDT-TS ∼90 for 10-residue fragments vs. GDT-TS ∼70 for whole proteins). This trend continues until the model is fully trained, where the gap between 10-residue fragments and whole proteins shrinks to a mere 10 GDT-TS points. Thus, while the model ultimately learns to predict global structure almost as well as local structure, it first learns to predict the latter.

**Figure 6:**
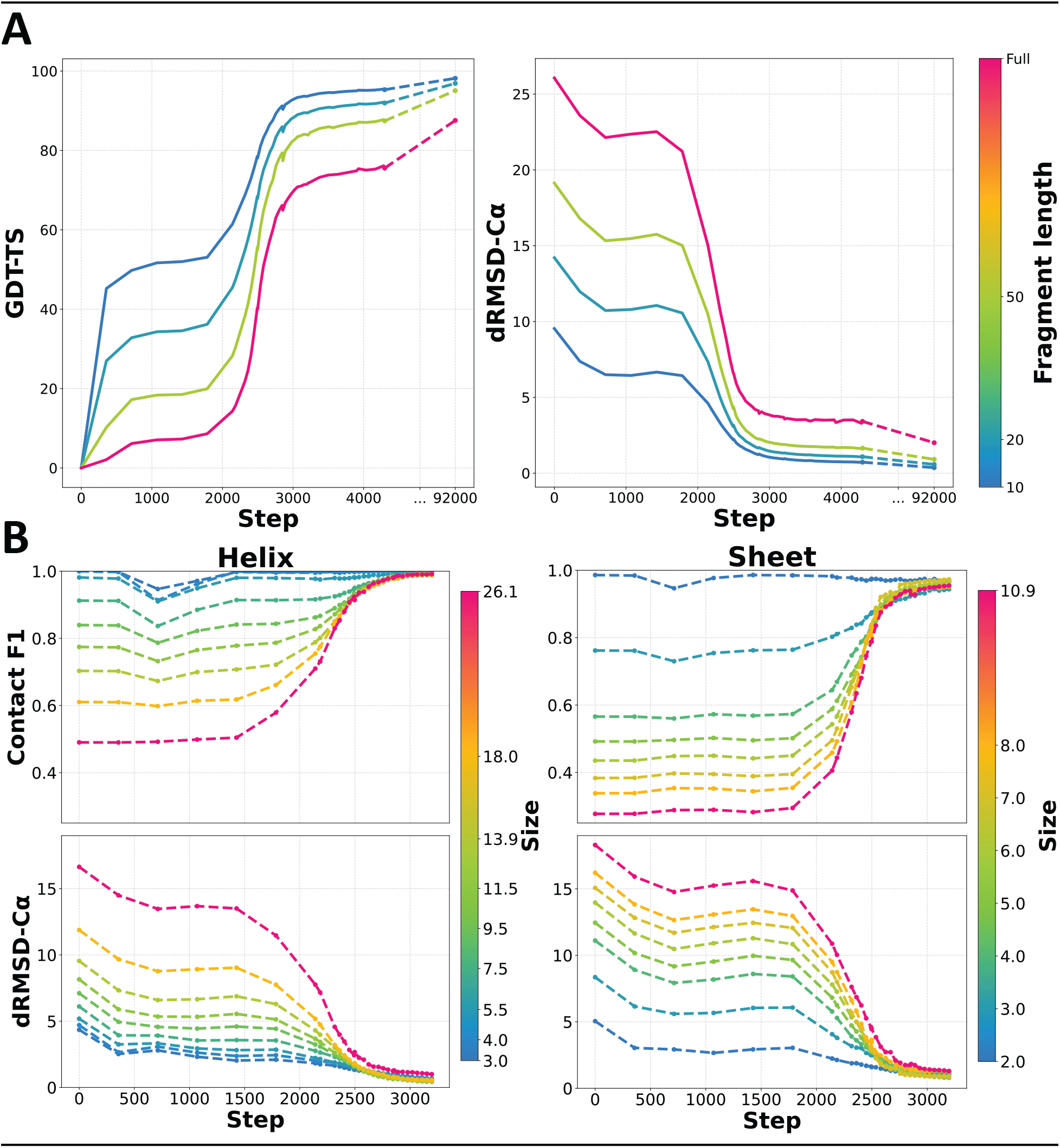
Learning proceeds at multiple scales. **(A)** Mean GDT-TS and dRMSD-Cα validation scores as a function of training step for non-overlapping protein fragments of varying lengths (colorbars indicate fragment length). **(B)** Average contact F1 score (8Å thresh-old) and dRMSD for predicted alpha helices and beta sheets of varying lengths and number of strands, respectively, as a function of training step. Colorbars indicate the weighted average of the lengths and widths of helices and sheets in each bin, respectively.

Turning to secondary structure, we investigated whether the same multi-scale learning behavior is detectable when examining SSEs. As before, we treat as ground truth the DSSP classifications of experimental structures in our validation set, focusing exclusively on alpha helices and beta sheets. We bin both SSEs according to size, defined as number of residues for alpha helices and number of strands for beta sheets. As uniform binnings would result in highly imbalanced bins, we instead opt for a dynamic binning procedure. First, each SSE is assigned to a (potentially imbalanced) bin that corresponds to its size. Bins are then iteratively merged with adjacent bins, subject to the condition that no bin exceed a maximum size (in this case, 200 for helices and 30 for sheets), until no further merges can occur. Finally bins below a minimum bin size (20 for both) are unconditionally merged with adjacent bins. We compute metrics averaged over each bin independently (Figure 6B).

Similar to what we observe for tertiary structure, short helices and narrow sheets are better predicted during earlier phases of training than their longer and wider counterparts, respectively. Improvement in SSE accuracy coincides with the rapid rise in tertiary structure accuracy, albeit shifted, as we observed in Figure 5A. Notably, the final quality of predicted SSEs is essentially independent of length/width despite the initially large spread in prediction accuracies, suggesting that OpenFold ultimately becomes scale-independent in its predictive capacity, at least for secondary structure. We note that the identification of SSEs is performed by DSSP, which is sensitive to the details of their hydrogen bonding networks. It is possible that in earlier phases of training, OpenFold has already recovered aspects of secondary structure not recognized by DSSP on account of imprecise atom positioning.

### OpenFold is more efficient and trains more stably than AlphaFold2

While the OpenFold model we used in all of the above experiments perfectly matches the computational logic of AlphaFold2, we have additionally implemented a number of changes that minimally alter model characteristics but improve ease of use and performance when training new models and performing large-scale predictions.

First, we made several improvements to the data preprocessing and training procedure, including a low-precision (“FP16”) training mode that facilitates model training on commercially available GPUs. Second, we introduced a change to the primary structural loss, FAPE, that enhances training stability. In the original model, FAPE is clamped—*i.e.,* limited to a fixed maximum value—in a large fraction of training batches. We find that in the dynamic early phase of training, this strategy is too aggressive, limiting the number of batches with useful training signal and often preventing timely convergence. Rather than clamping entire batches in this fashion, we instead clamp the equivalent fraction of samples within each batch, ensuring that each batch contains at least some unclamped chains. In doing so, we are able to substantially improve training stability and speed up model convergence (see Figure 7 and Appendix B.2).

**Figure 7:**
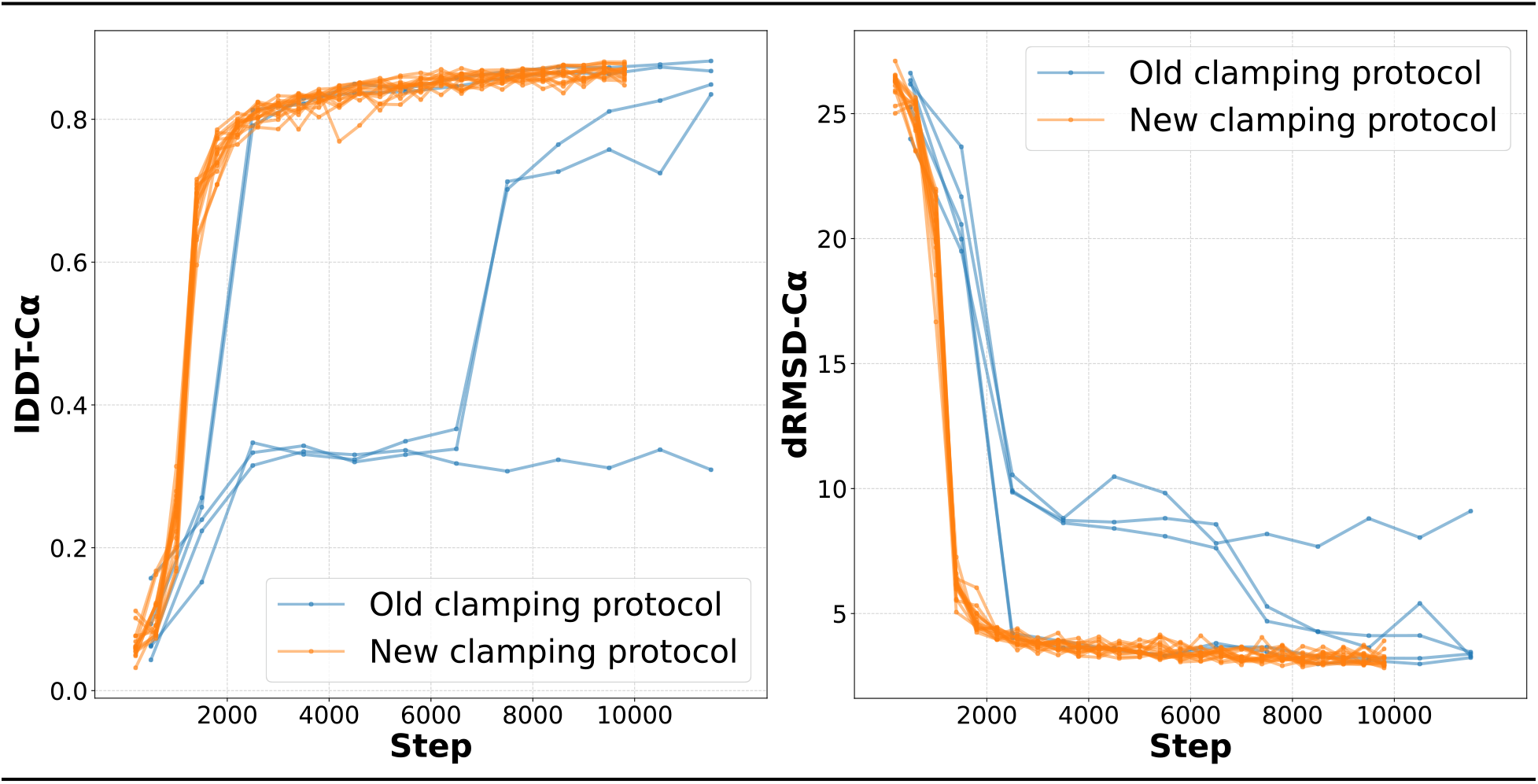
Stability of new FAPE clamping protocol. lDDT-Cα and dRMSD-Cα on CAMEO validation set as a function of training step for five independent training runs with (orange) and without (blue) new FAPE clamping protocol. Runs using old protocol exhibit substantial instability with two rapidly converging runs, two late converging runs, and one non-converging run. In contrast, all 15 independent runs using the new protocol converge rapidly. Runs using the new protocol also reach high accuracy faster.

Third, we made optimizations that improve memory efficiency during training, when model weights are continually updated to optimize model behavior for prediction, and inference, when the model is used to make new predictions. In AlphaFold2, the computational characteristics of these two modes vary greatly. To save memory at training time, which requires storing intermediate computations during the optimization procedure, AlphaFold2 and OpenFold are evaluated on short protein fragments ranging in size from 256 to 384 residues. At inference time, intermediate computations need not be stored, but input sequences can be more than ten times longer than the longest fragments encountered during training. Since the model’s memory usage naively grows cubically with input length, inference-time prediction stresses modules that are not necessarily bottlenecks at training time. To satisfy both sets of desiderata and enhance model efficiency, we implemented a number of training- and inference-specific optimizations. These optimizations create trade-offs between memory consumption and speed that can be tuned differently for training and inference. They include advanced implementations of neural network attention mechanisms ([44]) with favorable properties for unusually short and long sequences ([45], [29]), module refactoring for lower memory usage, optional approximations of certain computations that reduce the memory burden, and specialized low-level code customized for GPU hardware. For technical details see appendices E.1 and E.2.

Taken together, these optimizations result in a substantially more efficient implementation than AlphaFold2. We report OpenFold runtimes in Table 1. During inference, Open-Fold is up to three times faster than AlphaFold2 for proteins shorter than 1,100 residues. AlphaFold is faster than OpenFold for proteins between 1,100 and 2,400 residues in length, but thereafter AlphaFold2 crashes on single GPUs due to memory constraints. OpenFold runs successfully on longer proteins and complexes, with single-GPU predictions reaching up to 4,700 residues. OpenFold training speed matches or improves upon that of AlphaFold2, as reported by other researchers using OpenFold ([46], [47]).

**Table 1:** OpenFold vs. AlphaFold2 prediction speed. Prediction runtimes in seconds on a single A100 NVIDIA GPU for OpenFold and AlphaFold2 on proteins of varying lengths. OpenFold is faster than AlphaFold2 for proteins shorter than 1,100 residues and is able to predict longer proteins than AlphaFold2 on the same hardware.

| N | OpenFold (s) | AlphaFold (s) | Speedup |
| --- | --- | --- | --- |
| 100 | **2.8** | 8.8 | 3.14x |
| 200 | **6.7** | 13.9 | 2.07x |
| 300 | **13.0** | 21.9 | 1.68x |
| 400 | **22.0** | 33.7 | 1.53x |
| 500 | **34.7** | 50.8 | 1.46x |
| 600 | **50.8** | 72.8 | 1.43x |
| 700 | **71.0** | 102.5 | 1.44x |
| 800 | **96.4** | 135.4 | 1.40x |
| 900 | **136.5** | 177.1 | 1.30x |
| 1000 | **215.3** | 222.8 | 1.03x |
| 1100 | 288.7 | **278.1** | 0.96x |
| ... |  |  |  |
| 1500 | 614.1 | **549.1** | 0.89x |
| ... |  |  |  |
| 2000 | 1251.2 | **1081.1** | 0.86x |
| ... |  |  |  |
| 2500 | **2223.9** | OOM | ∞ |
| ... |  |  |  |
| 3000 | **3517.5** | OOM | ∞ |
| ... |  |  |  |
| 3500 | **6509.1** | OOM | ∞ |
| ... |  |  |  |
| 4000 | **10393.3** | OOM | ∞ |
| ... |  |  |  |
| 4500 | **12507.7** | OOM | ∞ |
| ... |  |  |  |
| 4700 | **13908.8** | OOM | ∞ |

## 3 Discussion

We have developed OpenFold, a complete open-source reimplementation of AlphaFold2 that includes training code and data. By training OpenFold from scratch and matching the accuracy of AlphaFold2, we have demonstrated the reproducibility of the AlphaFold2 model for protein structure prediction. Furthermore, the OpenFold implementation introduces technical advances over AlphaFold2, including markedly faster prediction speed. It is built using PyTorch, the most widely used deep learning framework, facilitating incorporation of OpenFold components in future machine learning models.

OpenFold immediately makes possible two broad areas of advances: (i) deeper analyses of the strengths, weaknesses, and learning behavior of AlphaFold2-like models and (ii) development of new (bio)molecular models that take advantage of AlphaFold2 modules. In this work, we have focused on the former. First, we assessed OpenFold’s capacity to learn from training sets substantially reduced in size. Remarkably, we found that even a 100-fold reduction in dataset size (0.76% models in Figure 2A) results in models more performant than the first version of AlphaFold. Stated differently, the architectural advances introduced in AlphaFold2 enable it to be 100x more data efficient than its predecessor, which at the time of its introduction set a new state of the art. These results demonstrate that architectural innovations can have a more profound impact on model accuracy than larger datasets, particularly in domains where data acquisition is costly or time-consuming, as is often the case in (bio)molecular systems. However, it merits noting that AlphaFold2 in general learns MSA-structure, not sequence-structure, relationships. MSAs implicitly encode a substantial amount of structural knowledge, as evidenced by early co-evolution-based structure prediction methods which were entirely unsupervised, making no use of experimental structural data ([48], [49]). Hence, the applicability of the AlphaFold2 architecture to problems that do not exhibit a co-evolutionary signal remains undemonstrated.

Our data elision results can be interpreted in light of recent work on large transformer-based language models that has revealed broadly applicable “scaling laws” that predict model accuracy as a simple function of model size, compute utilized, and training set size ([50], [51]). When not constrained by any one of these three pillars, models benefit from investments into the other two. These observations have largely focused on transformer-based architectures, of which AlphaFold2 is an example, but more recent work has revealed similar behavior for other architectures ([52]). Although determining the precise scaling properties of AlphaFold2 is beyond the scope of the present study, our results suggest that it is hardly constrained by the size or diversity of the PDB, motivating potential development of larger instantiations of its architecture.

By analyzing predicted structures of partially trained models, we discovered that AlphaFold2-like models learn spatial dimensions sequentially. This behavior has implications for the design of model architectures and training regimens. For example, integrating physical priors into machine learning models is an area of outstanding scientific interest ([53]). Efforts at such syntheses have had mixed results, and, indeed, AlphaFold2 serves as a seminal example of a highly successful model that is almost entirely devoid of physical priors. As we have mentioned, explicitly forbidding chemical violations—and, along with them, low-dimensional structures of the kind output by the model early in training—drastically alters AlphaFold2’s learning behavior. Our observation of the spatially collapsed learning phase provides an explanation for this observation. The solution that AlphaFold2 adopts for this problem, namely to penalize against physical violations only in later stages of training, suggests a broader strategy to tackle the incorporation of physical priors: a curriculum learning approach in which models are first free to extract information and learn from data, after which more complex physical priors can be gradually introduced to boost the model’s capacity for generalization. Analyzing learning trajectories, as we have done for OpenFold, provides a concrete timeline for when such priors can be injected into the training process. Finally, we observed that the spatially collapsed phases correspond to imperfect lower-dimensional PCA projections of the final predicted structure. Why this occurs is not *a priori* obvious, given that other end-to-end differentiable protein structure models do not exhibit the same behavior (see *e.g.,* [41]). Although we do not have direct evidence, we suspect that aspects of the AlphaFold2 architecture—specifically the FAPE loss function— likely drive this phenomenon. We speculate that the PCA-like progression allows the model to greedily minimize error by solving problems with the biggest payoff to the FAPE loss first, which by definition lie along the largest principal component of the ground-truth structure. Once solved, the model moves on to smaller problems lying along other, lower-dimensional projections. Were this to be the case, the staggering of spatial dimensions during learning would be contingent on the geometry of proteins in the training set. The extreme case of a training set composed entirely of long, slim, tubular proteins would produce even more dramatically staggered phases. Conversely, a training set composed of perfectly spherical proteins would exhibit even growth along all spatial dimensions. This behavior would be a function of the overall training set and would not necessarily get reflected in individual proteins. Regardless, these observations suggest that it may be possible to deliberately simplify other difficult problems in molecular modeling with a learning curriculum in which “toy” models are first trained to predict lower-dimensional projections of target molecules (or more generally, geometric objects) before being tasked to predict their fully realized instantiations.

OpenFold lays the groundwork for future efforts aimed at improving the AlphaFold2 architecture and repurposing it for new molecular modeling problems. Since the release of our codebase, there have been multiple efforts that either depend on OpenFold or directly build upon and extend it. These include the ESMFold method for protein structure prediction ([54]), which replaces MSAs with protein language models ([55], [56], [57]), and FastFold, a community effort that has implemented significant improvements including fast model-parallel training and inference ([46]). We expect future work to go further by disassembling OpenFold to attack problems beyond protein structure prediction. For instance, the evoformer module is a general-purpose primitive for reasoning over evolutionarily related sequences. DNA and RNA sequences also exhibit a co-evolutionary signal, with efforts aimed at predicting RNA structure from MSAs fast materializing (*e.g.*, [58], [59], [60]). It is plausible that even more basic questions in evolutionary biology, such as phylogenetic inference, may prove amenable to evoformer-like architectures. Similarly, AlphaFold2’s structure module, and in particular the invariant point attention mechanism, provide a general purpose approach for spatial reasoning over polymers, one that may be further extendable to arbitrary molecules. We anticipate that as protein structures and other biomolecules shift from being an output to be predicted to an input to be used, downstream tasks that rely on spatial reasoning capabilities will become increasingly important (*e.g.*, [61], [62]). We hope that OpenFold will play a key role in facilitating these developments.

## Supporting information

7RDT_A_animation

7LBU_A_animation

7DMF_A_animation

7B3A_A_animation

7DQ9_A_animation

## Code availability

OpenFold can be accessed at

https://github.com/aqlaboratory/openfold

It is available under the permissive Apache 2 Licence.

## Data availability

OpenProteinSet and OpenFold model parameters are hosted on the Registry of Open Data on AWS (RODA) and can be accessed at

https://registry.opendata.aws/openfold/

Both are available under the permissive CC BY 4.0 license.

## Author contributions

G.A. wrote and optimized the OpenFold codebase, generated data, trained the model, performed experiments, and maintained the GitHub repository. S.K. wrote data preprocessing code. G.A., N.B., and M.A. conceived of and managed the project, designed experiments, analyzed results, and wrote the manuscript. G.A., B.Z., Z.Z., N.Z., and A.N. ran ablations. All authors read and approved the manuscript. The Flatiron Institute provided compute for ablations, all data generation, and our main training experiments. NVIDIA performed training stability experiments, fixed critical bugs in the codebase, added new model features, and provided compute for ablations. StabilityAI provided compute for ablations.

## Acknowledgements

We would like to thank the Flatiron Institute, OpenBioML, Stability AI, the Texas Advanced Computing Center, and NVIDIA for providing compute for experiments in this paper. Individually, we would like to thank Milot Mirdita, Martin Steinegger, and Sergey Ovchinnikov for valuable support and expertise. This research used resources of the National Energy Research Scientific Computing Center, which is supported by the Office of Science of the U.S. Department of Energy under Contract No. DE-AC02-05CH11231. The authors acknowledge the Texas Advanced Computing Center (TACC) at The University of Texas at Austin for providing HPC resources that have contributed to the research results reported within this paper. N.B. is supported by DARPA PANACEA program grant HR0011-19-2-0022 and NCI grant U54-CA225088. B.Z. and Z.Z. are supported by grants NSF OAC-2112606 and OAC-2106661.

## Appendix

### A CASP15 results

While OpenFold did participate in CASP15, model training was not completed until late June 2022, more than a month after the competition began. Until the final weeks of evaluation, OpenFold’s CASP15 predictions were also adversely affected by an error in its inference pipeline. In Supplementary Figure 1, we plot the accuracy of predictions on CASP15 domains by both AlphaFold2 and the finalized OpenFold.

**Supplementary Figure 1:**
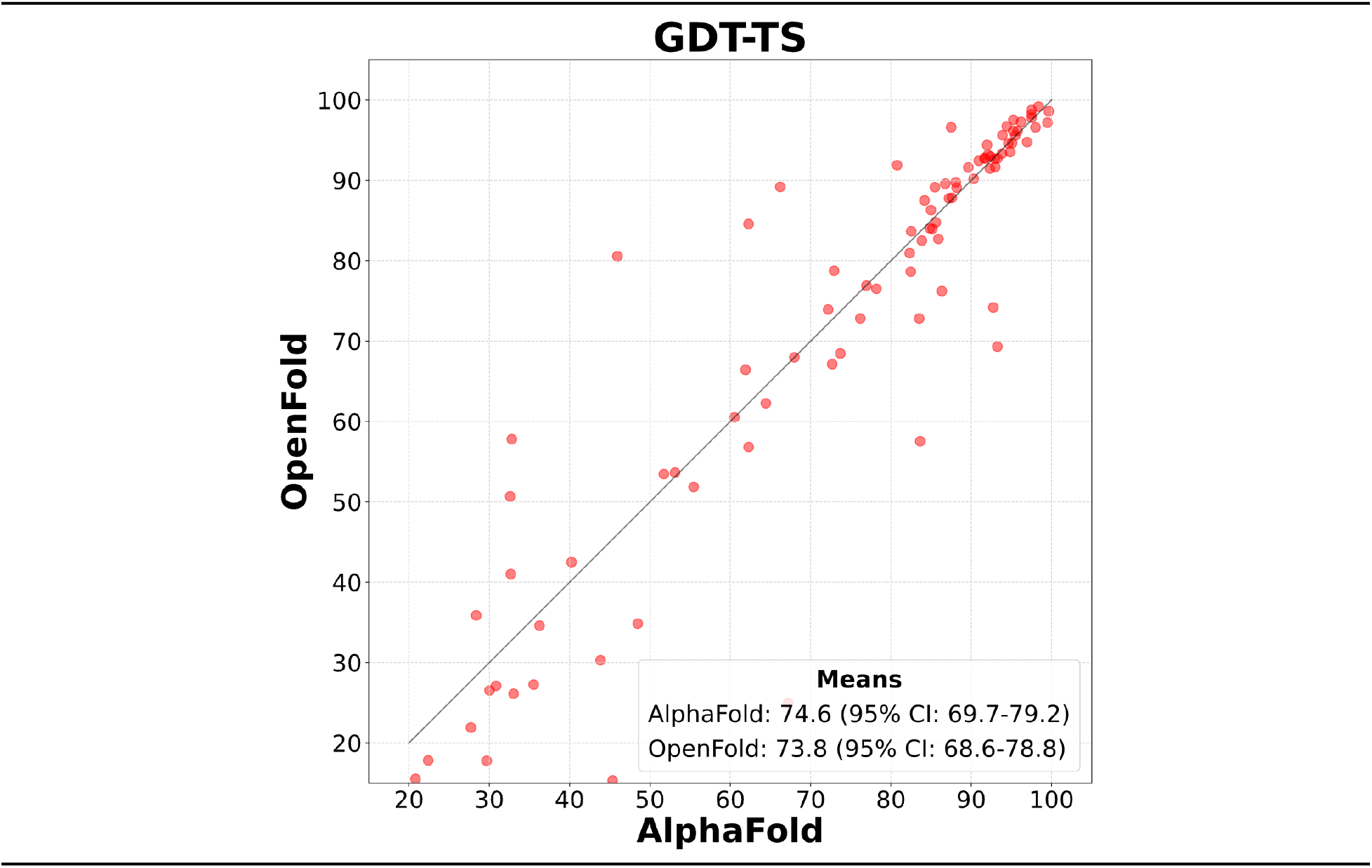
OpenFold matches the accuracy of AlphaFold2 on CASP15 targets. Scatter plot of GDT-TS values of AlphaFold and OpenFold “Model 1” predictions against all currently available “all groups” CASP15 domains (n = 90). OpenFold’s mean accuracy (95% confidence interval = 68.6-78.8) is on par with AlphaFold’s (95% confidence interval = 69.7-79.2) and OpenFold does at least as well as the latter on exactly 50% of targets. Confidence intervals of each mean are estimated from 10,000 bootstrap samples.

### B Differences between OpenFold and AlphaFold2

In this section we describe additions and improvements we made to OpenFold subject to the constraint that the weights of the two models should be interchangeable. We also describe our design decisions in the handful of cases where the AlphaFold2 paper was ambiguous.

#### B.1 Changes to the data pipeline

##### Template trick

During AlphaFold2 training, structural template hits undergo two successive rounds of filtering. Between these two rounds, the dataloader parses the template structure data. The top 20 template hits to pass both filters are shuffled uniformly at random. Finally, the dataloader samples a number of templates uniformly at random in the range and draws that many samples from the shuffled pool of valid hits. These are then passed as inputs to the model. This subsampling process is intended to lower the average quality of templates seen by the model during training. We note that preemptively parsing structure files for each template hit during the filtering process is an expensive operation and, for proteins with many hits, considerably slows training. For this reason, we replace the original algorithm with an approximation. Instead of sampling hits from the top 20 template candidates from the pool of templates that pass both sets of filters, we use the top 20 hits to pass the first filter. These are then shuffled and subsampled as before. Only when a hit is drawn is it passed through the second filter; hits that fail to pass the second filter at this point are discarded and replaced. If not enough hits in the initial 20-sample pool pass the second filter, we continue drawing candidates from the top hits outside that pool, without further shuffling. This procedure has the disadvantage that, if too many hits pass the first filter but not the second, the hits used for the model are not shuffled. Even in cases where only of the initial hits fail to pass the second filter, OpenFold effectively only shuffles the top proteins to pass both filters, strictly increasing the expected quality of template hits relative to those used by AlphaFold2. However, in most cases, this approximation allows the dataloader to parse only as many structure files as are needed, speeding up the process by a factor of at least 5. In practice, the vast majority of invalid template hits are successfully detected by the first filter, suggesting that the difference in final template quality between the two procedures is marginal.

##### Self-distillation training set filtering

The 270,262 MSAs yielded by our self-distillation procedure is smaller than the 355,993 reported by DeepMind, despite having started with the same database. We suspect that the discrepancy arises due to the first step of the filtering process, of which the description in the AlphaFold2 paper is somewhat ambiguous.

##### Zero-centering target structures

We find that centering target structures at the origin slightly improves the numerical stability of the model, especially during low-precision training.

#### B.2 Plateaus and phase transitions

During training of the original version of OpenFold, we and third party developers observed two distinct training behaviors. The large majority of training runs are almost identical to the training curves shown in Figure 1; after a few thousand training steps, validation lDDT-C rapidly rises to ∼0.83 and improves only incrementally thereafter. Occasionally, such runs exhibit a “double descent,” briefly improving and then degrading in accuracy before finally converging in the same way. In a fraction of training runs, however, lDDT-C plateaus between 0.30 and 0.35 on the same set (see Figure 7A). Anecdotally, these values appear to be consistent across environments, OpenFold versions, architectural modifications, and users. If the runs are allowed to continue long past the point where lDDT-C would otherwise have stopped improving (>10k training steps), they eventually undergo a phase transition, suddenly exceeding 0.8 lDDT-C and then continuing to improve much as normal runs do. We have not been able to determine whether this phenomenon is the result of an error in the OpenFold codebase or if it is a property of the AlphaFold2 algorithm.

Since running the experiments described in the main text of this paper, we have discovered a workaround that deviates slightly from the original AlphaFold2 training procedure but that appears to completely resolve early training instabilities. In the original training configuration, for each AlphaFold2 batch, backbone FAPE loss, the model’s primary structural loss, is clamped for all samples in the batch with probability 0.9. This practice is potentially problematic during the volatile early phase of training, when FAPE values can be extremely large and frequent clamping zeroes gradients for most of the residues in each crop. We find that clamping each sample independently, *i.e.,* clamping approximately 90% of the samples in each batch rather than clamping 90% of all batches, eliminates training instability and speeds up convergence to high accuracy by about 30%. We show before-and-after data in Figure 7B.

### C Training details

Our main OpenFold model was trained using the abridged training schedule outlined in Table 4 of the AlphaFold2 Supplementary Materials rather than the original training schedule in Table 5. Specifically, it was trained for three rounds: the initial training phase, the fine-tuning phase, and the predicted-TM ([63]) fine-tuning stage. During the initial training phase, sequences were cropped to 256 residues, MSA depth was capped at 128, and extra MSA depth was capped at 1,024. During fine-tuning, these values were increased to 384, 512, and 5,120, respectively. The second phase also introduced the “violation” and “experimentally resolved” losses, which respectively penalize non-physical steric clashes and incorrect predictions of whether atomic coordinates are resolved in experimental structures. Next, we ran a short third phase with the predicted TM score loss enabled. The three phases were run for 10 million, 1.5 million, and 0.5 million protein samples, respectively. We trained the model with PyTorch v1.10, DeepSpeed ([26]) v0.5.10, and stage 2 of the ZeRO redundancy optimizer ([64]). We used Adam ([65]) with _1_, _2_, and ^−6^. We warmed up the learning rate linearly over the first 1,000 iterations from 0 to 10^−3^. After approximately 7 million samples, we marginally decreased the learning rate to ^−4^. This decrease had no noticeable effect on model training. For the latter two phases, the learning rate was halved to ^−4^. All model, data, and loss-related hyperparameters were identical to those used during AlphaFold2 training. We also replicated all of the stochastic training-time dataset augmentation, filtering, and resampling procedures described in the original paper.

During the initial fine-tuning and subsequent predicted-TM fine-tuning phases, we manually sampled checkpoints at peaks in the validation lDDT-C ([33]). These checkpoints were added to the pool of model checkpoints used in the final model ensemble.

Training was run on a cluster of 44 NVIDIA A100 GPUs, each with 40GB of DRAM. The model was trained in a data-parallel fashion, with one protein per GPU. In order to simulate as closely as possible the batch size of 128 used in training AlphaFold2, we performed three-way gradient accumulation to raise our effective batch size from 44 to 132. Supplementary Figure 2 contains additional data from the training run.

As in the original paper, CAMEO chains longer than 700 residues were removed from the validation set.

**Supplementary Figure 2:**
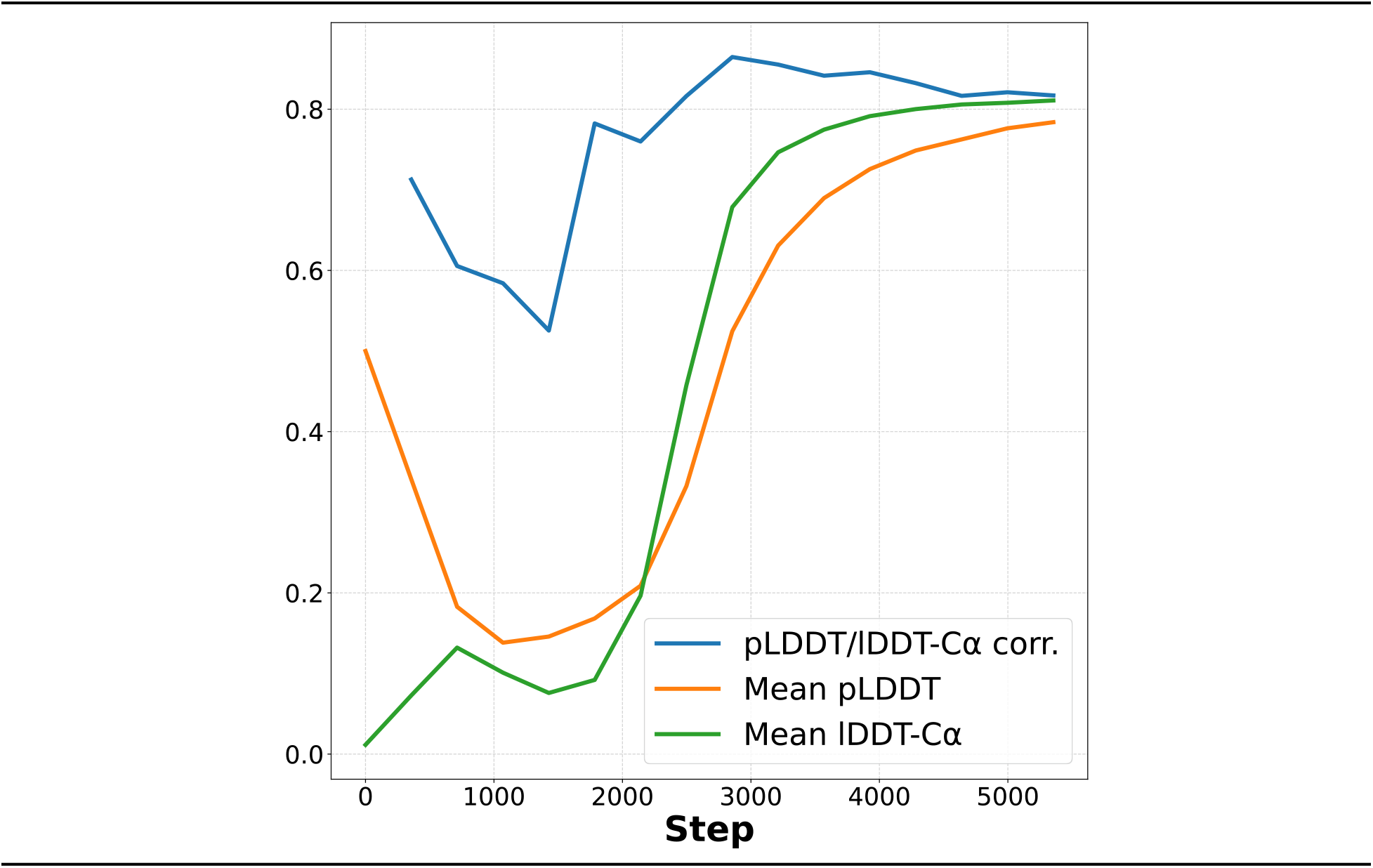
Mean correlation between lDDT-Cα and pLDDT over validation set chains as a function of training step in the early stage of training. Mean values for both metrics are superimposed.

### D Inference details

Runtime benchmarks were performed on a single 40GB A100 GPU. Times correspond to the intensive ‘model_1_ptm’ config preset, which uses deep MSAs and the maximum number of templates.

For proteins shorter than 1,000 residues, we take advantage of OpenFold’s TorchScript tracing capability and FlashAttention. For longer proteins, both tools become unstable and so we disable them. We plan to address this shortcoming in the future.

For runtime benchmarks we upgraded to PyTorch v1.12 for its improvements to TorchScript. AlphaFold was run with JAX v. 0.3.13.

For reference, we include the distribution of lengths of our 132,000 PDB chains in Supplementary Figure 3.

**Supplementary Figure 3:**
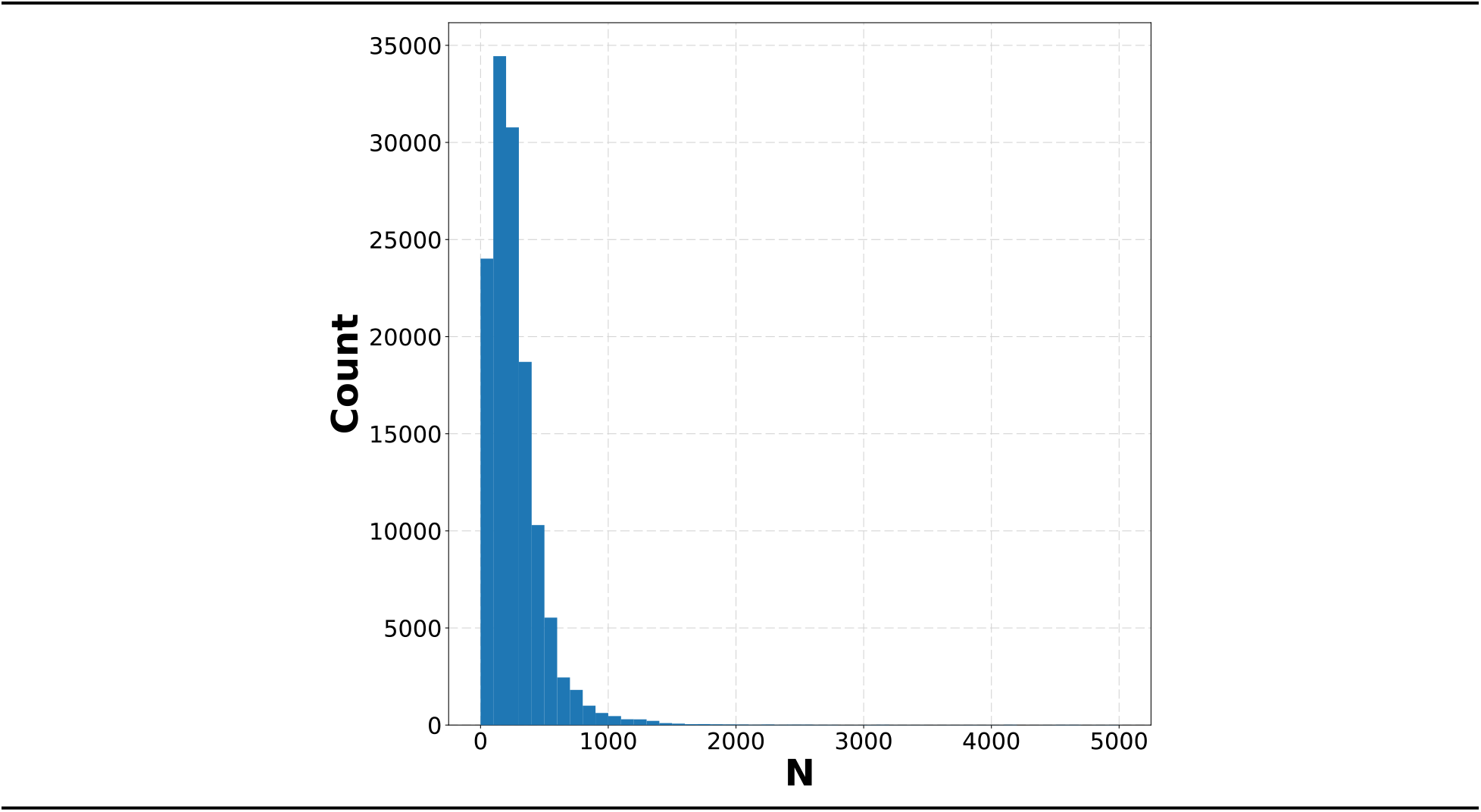
PDB sequence lengths are heavily concentrated in the region where OpenFold has an inference-time advantage. Binned sequence lengths of the 132,000 chains in the PDB training set.

### E Additional model optimizations and features

#### E.1 Training-time optimizations and features

Despite its relatively small parameter count (∼93M), AlphaFold2 manifests very large intermediate activations during training, resulting in peak memory usage—along with floating point operation counts—much greater than that of state-of-the-art transformer-based models from other domains ([66]). In AlphaFold2, peak memory usage during training grows cubically as a function of input sequence length. As a result, during the second phase of training when the inputs are longest, the model manifests individual tensors as large as 12GB. Intermediate activation tensors stored for the backward pass are even larger. This bottleneck is exacerbated by several limitations of the PyTorch framework. First, PyTorch is run eagerly and doesn’t benefit from the efficient compiler used by JAX models that improves runtime and reduces memory usage. Second, even on GPUs that in principle have sufficient memory to store all intermediate tensors used during the forward pass, suboptimal allocation patterns frequently result in memory fragmentation, preventing the model from utilizing all available memory. For this reason, among others, a preliminary version of OpenFold naively modeled after the official JAX-based implementation frequently ran out of memory despite having allocated as little as 40% of total available memory. To ameloriate these problems, we introduced several features that reduce peak memory consumption during training.

##### In-place operations

We refactored the model by replacing element-wise tensor operations with in-place equivalents wherever possible to prevent unnecessary allocation of large intermediate tensors.

##### Custom CUDA kernels

We implemented custom CUDA kernels for the model’s “MSA row attention” module, the multi-head attention operation where the aforementioned 12GB tensor is allocated. Modified from optimized softmax kernels from FastFold ([46]), which are in turn derived from OneFlow kernels ([67]), our kernels operate entirely in-place. This is made possible partly by a fusion of the backward passes of the softmax operation and the succeeding matrix multiplication. Overall, only a single copy of the quadratic attention logit tensor is allocated, resulting in peak memory usage 5 and 4 times lower than equivalent native PyTorch code and the original FastFold kernels, respectively.

##### DeepSpeed

OpenFold is trained using DeepSpeed. Using its ZeRO Redundancy optimizer ([64]) in the “stage 2” configuration, the model partitions gradients and optimizer states between GPUs during data-parallel training, further reducing peak memory usage.

##### Half-precision training

By default, as a memory-saving measure, AlphaFold2 is trained using bfloat16 floating point precision. This 16-bit format trades the large precision of the classic half-precision format (FP16) for the complete numerical range of full-precision floats (FP32), making it well-suited for training deep neural networks of the type used by AlphaFold2, which is not compatible with FP16 training by default. However, unlike FP16, bfloat16 hardware support is still limited to relatively recent NVIDIA GPUs (Ampere and Hopper architectures), and so the format remains out of reach for academic labs with access to older GPUs that are otherwise capable of training AlphaFold2 models (*e.g.,* V100 GPUs). We address this problem by implementing a stable FP16 training mode with more careful typecasting throughout the model pipeline, making OpenFold training broadly accessible.

#### E.2 Inference

We also introduce several inference-time optimizations to OpenFold. As previously mentioned, these features trade off memory usage for runtime, contributing to more versatile inference on chains of diverse lengths.

##### FlashAttention

We incorporate FlashAttention ([29]), an efficient fused attention implementation that tiles computation in order to reduce data movement between different levels of GPU memory, greatly improving peak memory usage and runtime in the process. We find it to be particularly effective for short sequences with 1,000 residues or less, on which it contributes to an OpenFold speedup of up to 15% despite only being compatible with a small number of the attention modules in the network.

##### Low-memory attention

Separately, OpenFold makes use of a recent attention algorithm that uses a novel chunking technique to perform the entire operation in constant space ([45]). Although enabling this feature marginally slows down the model, it nullifies attention as a memory bottleneck during inference.

##### Refactored triangle multiplicative attention

A naive implementation of the triangle multiplicative update manifests 5 concurrent tensors the size of the input pair representation. These pair representations grow quadratically with input length, such that during inference on long sequences or complexes, they become the key bottleneck. We refactored the operation to reduce its peak memory usage by 50%, requiring just 2.5 copies of the pair representation.

##### Template averaging

AlphaFold2/OpenFold create separate pair embeddings for each structural template passed to each model, then reduce them to a single embedding at the end of the template pipeline with an attention module. For very long sequences, or very many templates, this operation can become a memory bottleneck. AlphaFold-Multimer ([22]) avoids this problem by computing a running average of template pair embeddings. Although we trained OpenFold using the original AlphaFold2 (non-multimer) procedure, we find that the newer approach can be adopted during inference without a noticeable decrease in accuracy. We thus make it available as an optional inference-time memory-saving optimization.

##### In-place operations

Without the requirement to store intermediate activations for the backward pass, OpenFold is able to make more extensive use of in-place operations during inference. We also actively remove unused tensors to mitigate crashes caused by memory fragmentation.

##### Chunk size tuning

AlphaFold2 offsets extreme inference-time memory costs with a technique called “chunking,” which splits input tensors into “chunks” along designated, module-specific sub-batch dimensions then runs those modules sequentially on each chunk. In AlphaFold2’s case, the chunk size used in this procedure is a model-wide hyperparameter that is manually tuned. OpenFold, on the other hand, dynamically adjusts chunk size values for each module independently, taking into account the model’s configuration and the current memory limitations of the system. Although the profiling runs introduced by this process incur a small computational overhead, the modules do not need to be recompiled for each run, unlike their AlphaFold2 equivalents, and said profiling runs are only necessary the first time the model is run; once computed, the optimal chunk sizes are cached and reused until conditions change. We find this to be a robust way to seamlessly improve runtimes in a variety of settings.

##### Tensor offloading

Optionally, OpenFold can aggressively offload intermediate tensors to CPU memory, temporarily freeing additional GPU memory for memory-intensive computations at the cost of a considerable slowdown. This feature is useful during inference on extremely long sequences that would otherwise not be computable.

##### TorchScript tracing

Specially written PyTorch programs can be converted to TorchScript, a JIT-compiled variant of PyTorch. We use this feature during inference to speed up parts of the Evoformer module. Although TorchScript tracing and compilation do introduce some overhead at the beginning of model inference, and lock the model to a particular sequence length similar to JAX compilation, we find that using TorchScript achieves overall speedups of up to 15%, especially on sequences shorter than 1,000 residues. This feature is particularly useful during batch inference, where sequences are grouped by length to avoid repeated re-compilations and take maximal advantage of faster inference times.

**Supplementary Figure 4:**
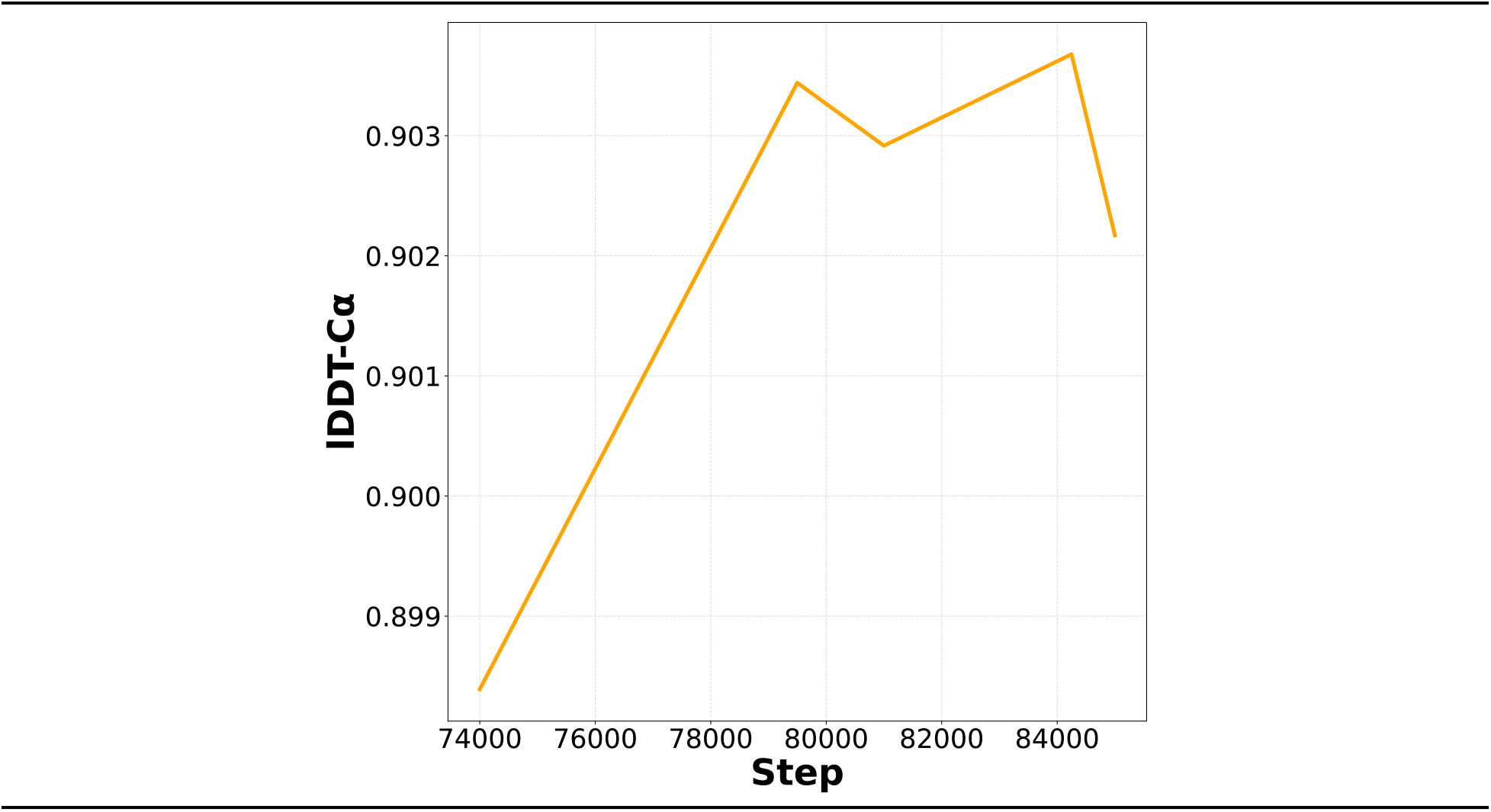
Fine-tuning does not materially improve prediction accuracy on long proteins. Mean lDDT-C over validation proteins with at least 500 residues as a function of fine-tuning step.

##### AlphaFold-Gap implementation

OpenFold currently supports multimeric inference using AlphaFold-Gap ([13]), a zero-shot hack that allows inference on protein complexes using monomeric weights. Although it falls short of the accuracy of AlphaFold-Multimer^4^ it is a capable tool, especially for homomultimers. Since complexes manifest in the model as long sequences, OpenFold-Gap in particular benefits from the memory optimizations discussed earlier.

### F Extended analysis

#### F.1 Effect of fine-tuning on long proteins

In Supplementary Figure 4, we illustrate the effect of fine-tuning on the lDDT-Cα of long chains. Though we observe a larger increase here than is seen for all proteins in Figure 1D, it is still less than half a point.

**Supplementary Figure 5:**
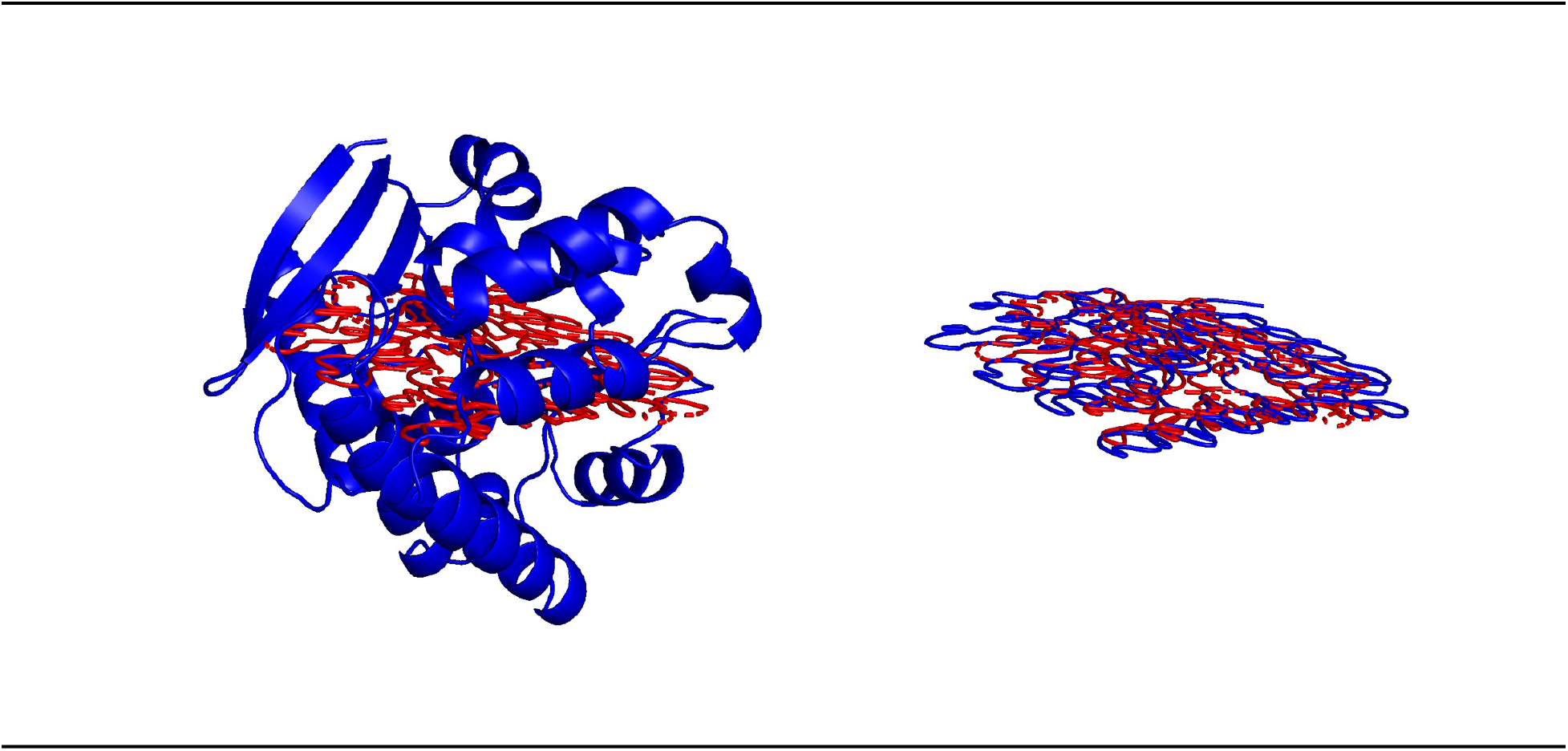
A “2D vs. 3D” comparison (left, corresponding to Figure 4A) and a “2D vs. 2D” comparison (right, corresponding to Figure 4B). The blue, two-dimensional structure on the right is the 2D PCA projection of the 3D structure on the left. The red structure in both images is the same 2D PCA projection of a prediction from the two-dimensional phase.

#### F.2 Dimensionality of output structures

Here we provide additional analyses of the staged learning of dimensionality. First, supplementary Figure 5 provides a visual aid for the projection operations used to produce Figure 4A and 4B. Second, because the precise timing of the phases of dimensionality differ slightly for individual proteins, we include for reference Supplementary Figure 6, which shows individual eigenvalues for all proteins in the validation set.

Third, for an additional perspective on the low-dimensionality phenomenon, we consider the radius of gyration, a popular measure of the compactness of a structure ([68]), given by

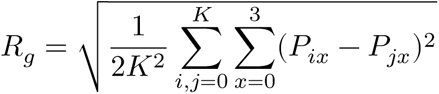

where *K* is the number of atoms in the protein and *P* ∈ ℝ^*K*×3^ is a coordinate vector. Real proteins obey known protein-phase-specific radius scaling laws (see e.g. [69]), and we wish to determine exactly how and when OpenFold begins to produce plausible structures that do the same. To accomplish this, we compute the radius of gyration of each of the experimental structures in our validation set and compare them to the radii of gyration of model predictions as a function of training step. Results are shown in Supplementary Figure 7.

**Supplementary Figure 6:**
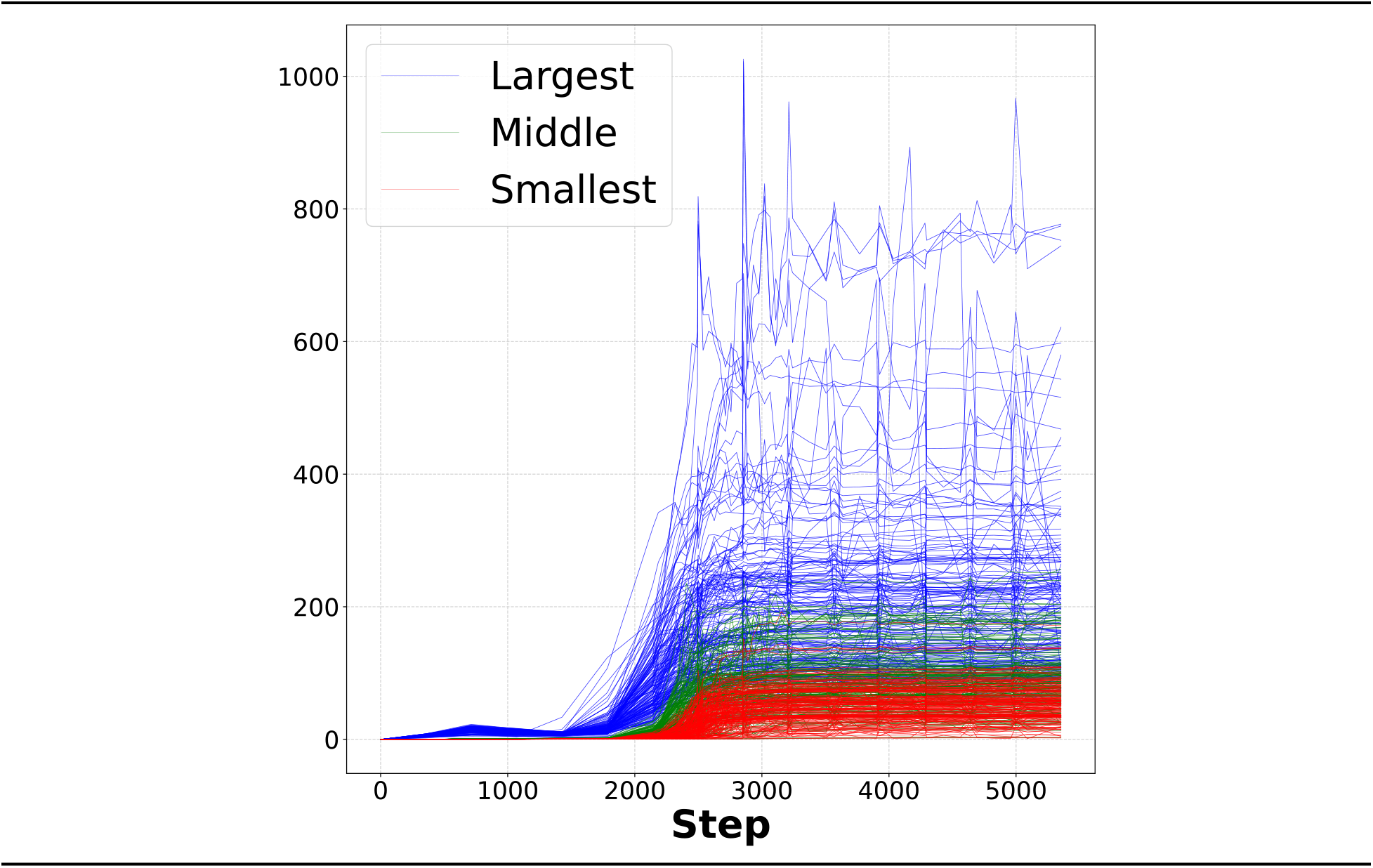
Sorted PCA eigenvalues for all proteins in the CAMEO validation set as a function of OpenFold training step.

The plots correspond approximately to the phases of dimensionality illustrated in 4. In the first two panels, before the model enters the three-dimensional phase, the radii of gyration of predicted structures are indeed systematically smaller than those of experimental structures. Immediately thereafter, the radii of gyration are largely correct, and only minor adjustments are made in the final panel.

#### F.3 DSSP state reduction

We reduce the 8-state DSSP assignment to 3 states using the following mapping:

**Supplementary Figure 7:**
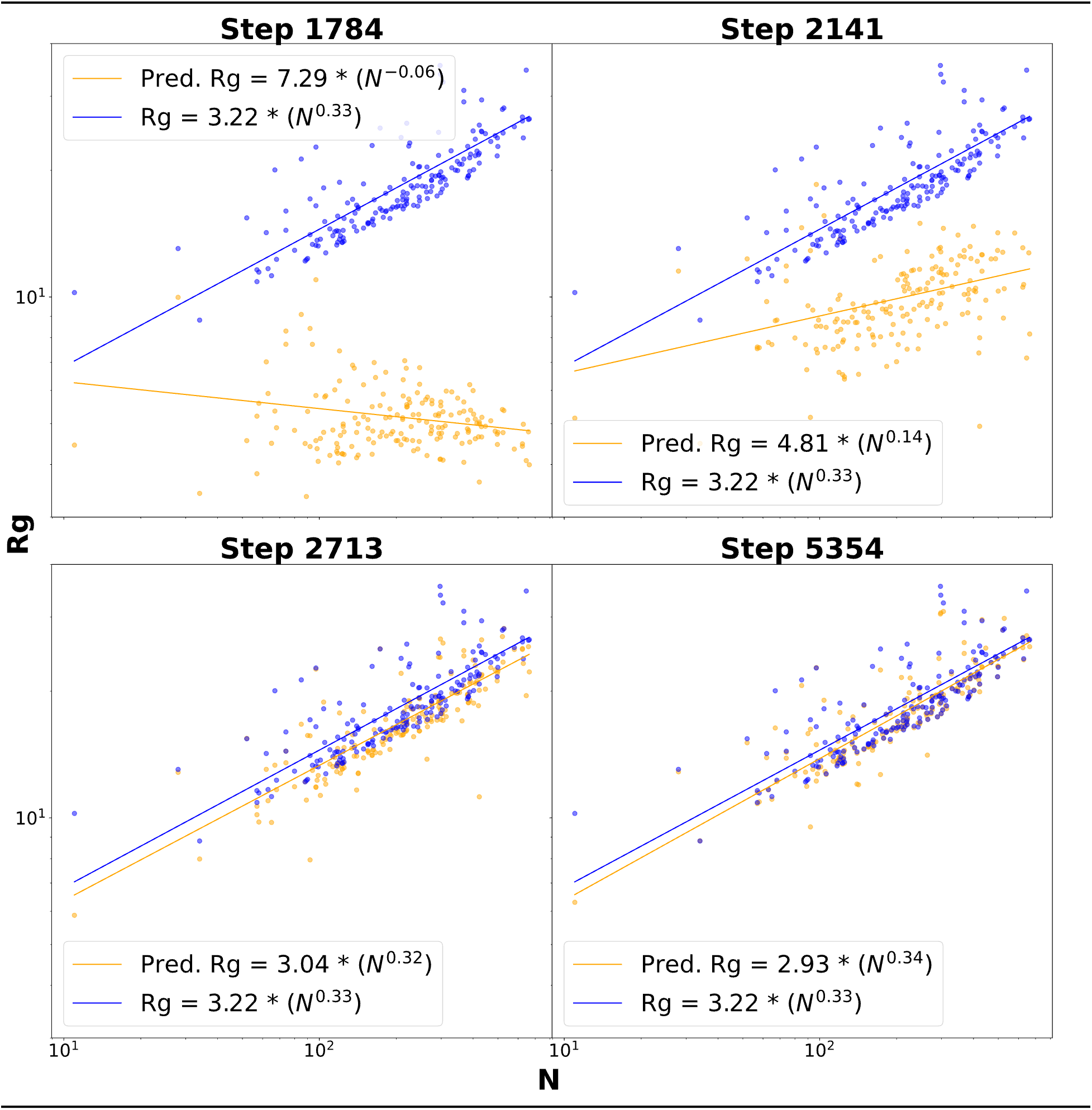
Radius of gyration as an order parameter for learning protein phase structure. Radii of gyration for proteins in the CAMEO validation set (orange) as a function of sequence length over training time, plotted on a log-log scale against experimental structures (blue). Legends show equations of best fit curves, computed using non-linear least squares. The training steps chosen correspond loosely to the four phases of dimensional growth.

**Supplementary Figure 8:**
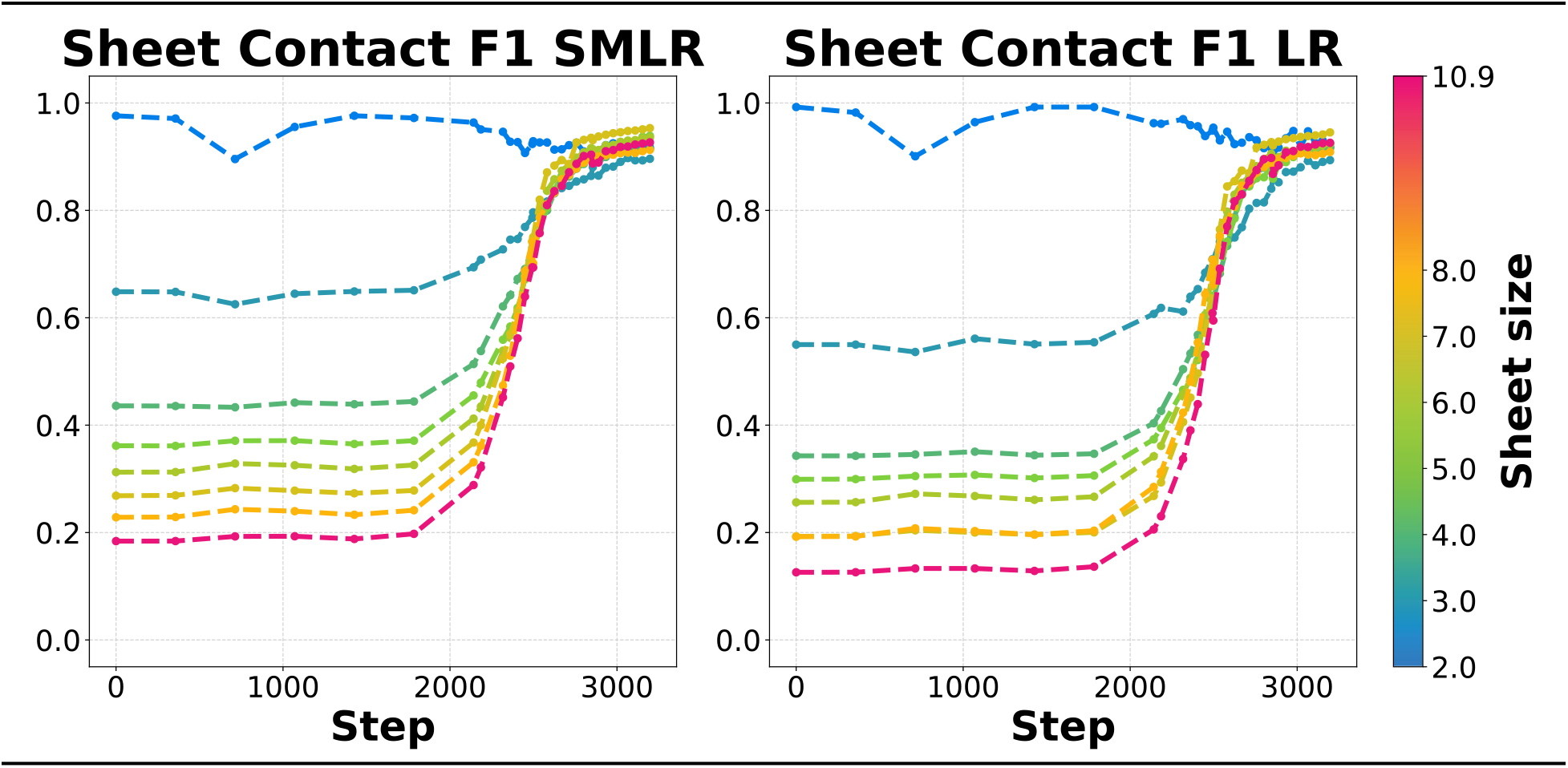
Contact prediction for beta sheets at different ranges. Binned contact F1 scores (8Å threshold) for beta sheets of various widths as a function of training step at different residue-residue separation ranges (SMLR ≥ 6 residues apart; LR ≥ 24 residues apart, as in [8]). Sheet widths are weighted averages of sheet thread counts within each bin, as in Figure 6B.

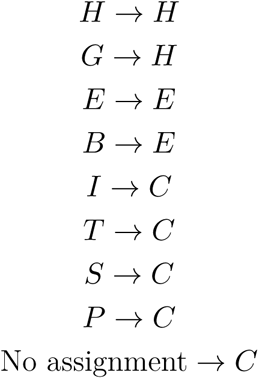

where ‘C’ denotes “coil” and ‘E’ denotes “strand”.

#### F.4 Additional secondary structure data

In Supplementary Figure 8, we provide another view of the sheet data in Figure 6B by distinguishing between small- (S: 6Å), medium- (M: 12Å) and long- (L: 24Å) range contacts, as in *e.g.*, ([8]).

**Supplementary Table 1:** Secondary structure recovery by class-stratified models. Recall and F1 scores for reduced secondary structure categories derived using DSSP. Results are shown for the two class-stratified models from the final panel of Figure 2B, here evaluated on the CAMEO validation set. The reduced secondary state scheme described in Appendix F.3 is used.

| Ablated CATH category | Training set | Mean CAMEO lDDT-Cα |
| --- | --- | --- |
| (T)opology | 100% avail. T | 0.806 |
|  | 50% avail. T | 0.786 |
|  | 10% avail. T | 0.678 |
|  | 5% avail. T | 0.567 |
| (A)rchitecture | 100% avail. A | 0.795 |
|  | 50% avail. A | 0.763 |
|  | 10% avail. A | 0.627 |
|  | 5% avail. A | 0.586 |
| (C)lass | Class 1 (“Mostly alpha”) | 0.689 |
|  | Class 2 (“Mostly beta”) | 0.713 |

#### F.5 Data elision validation using CAMEO

In order to properly assess the model’s generalization capacity, we evaluated each set of CATH ablations on a corresponding validation set in the main text. *I.e.,* the T ablations were evaluated on held-out topologies, the A ablations on held-out architectures, and so on. As a result, different data elision experiments cannot not be compared directly. For a more consistent picture of the relative final accuracies of each set of data elision experiments, we reevaluate the final checkpoints of each model on our standard CAMEO validation set in Supplementary Table 1.

Note that a potential confounding factor is that CATH classifications are not yet available for proteins in the CAMEO validation set, making it difficult to determine the degree of overlap in fold space between the training set of each data elision and the validation set. If the CAMEO validation set happens to contain chains with architectures in the training sets of the smaller A ablations, for example, the values in Supplementary Table 1 would overestimate the accuracies of the corresponding models.

#### F.6 Secondary structure recovery of class-stratified models

In Figure 2C, we show predictions of two class-stratified models for two CAMEO chains. For a more comprehensive picture, we report mean reduced-state DSSP recall and F1 over the entire CAMEO validation set for both models in Supplementary Table 2.

#### F.7 Characteristics of class-stratified training sets

We note in the main text that domains in the class used to train the “Mainly alpha” class elision still contain some beta sheets, and vice versa. To quantify this, in Supplementary Figure 9 we show the distribution of alpha helices and beta sheets of different sizes in the two class elision training sets based on 1,000 randomly chosen samples.

**Supplementary Figure 9:**
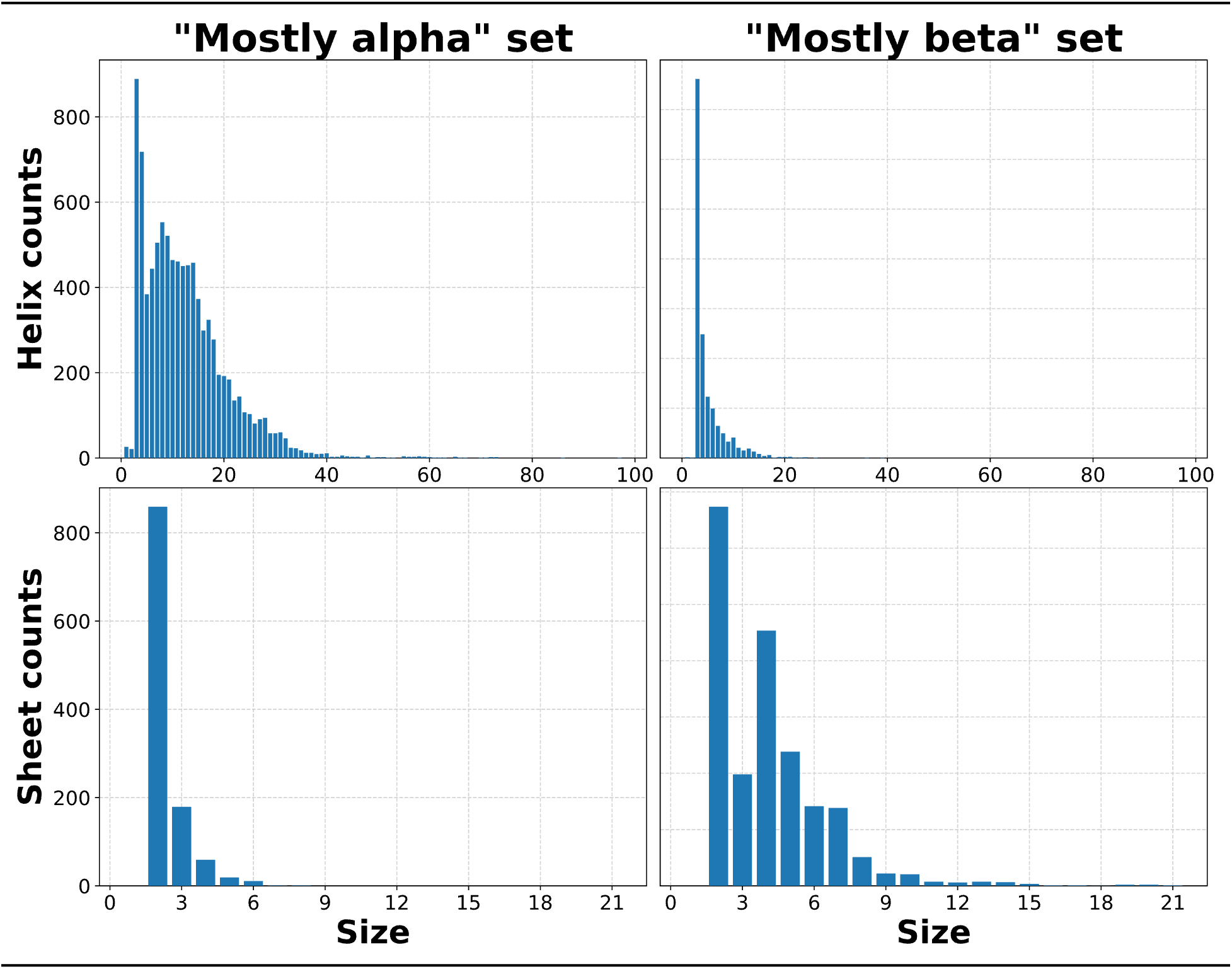
The “Mostly alpha” CATH class contains some beta sheets, and vice versa. Counts for alpha helices and beta sheets in the mostly alpha and mostly beta CATH class-stratified training sets from Figure 2, based on 1,000 random samples. Counts are binned by size, defined as the number of residues for alpha helices and number of strands for beta sheets.

**Supplementary Table 1:**
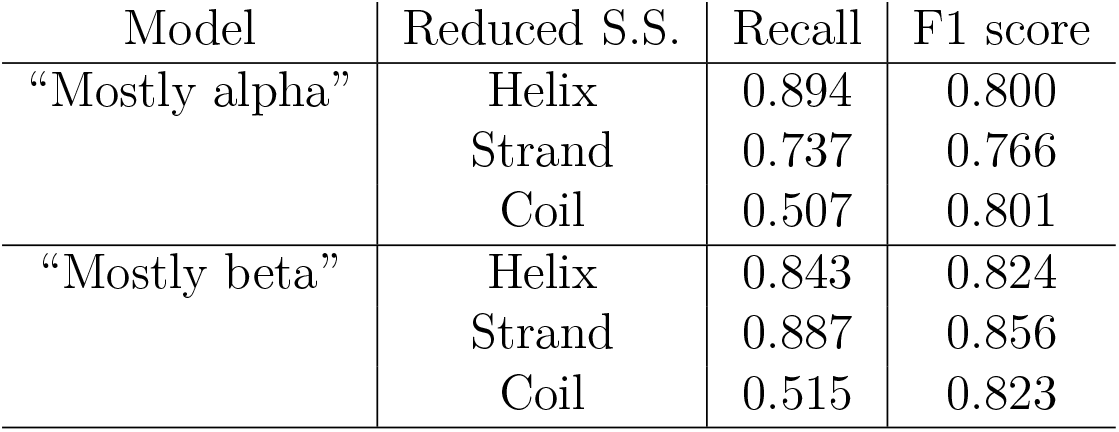
Data elision models evaluated on CAMEO validation set. Rows correspond to CATH elisions reported in Figure 2, except evaluations reported here are based on the CAMEO validation set.

### G Known issues during training

During and after the primary OpenFold retraining experiment, we discovered a handful of minor implementation errors that, given the prohibitive cost of retraining a full model from scratch, could not be corrected. In this section, we describe these errata and the measures that we have taken to mitigate them.

#### G.1 Distillation template error

As described in the main text, OpenFold/AlphaFold2 training consists of three phases, of which the first is the longest and most determinative of final model accuracy. During this first phase of the main OpenFold training run, a bug in the dataloader caused distillation templates to be filtered entirely; OpenFold was only presented with templates for PDB chains, which constitute ∼25% of training samples, and not self-distillation set chains. The issue was corrected for later phases, which were run slightly longer than usual to compensate. Although the accuracy of the resulting OpenFold model matches that of the original AlphaFold2 in holistic evaluations, certain downstream tasks that specifically exploit the template stack ([70]) do not perform as well as the original AlphaFold2. There is evidence, for instance, that OpenFold disregards the amino acid sequence of input templates. After the bug was corrected, follow-up experiments involving shorter training runs showed template usage behavior at parity with AlphaFold2. Furthermore, OpenFold can be run with the original AlphaFold2 weights in cases where templates are expected to be important to take advantage of the new inference characteristics without diminution of template-related performance.

#### G.2 Gradient clipping

OpenFold, unlike AlphaFold2, was trained using per-batch as opposed to per-sample gradient clipping (first noted by the UniFold team ([47])). UniFold experiments show that models trained using the latter clipping technique achieve slightly better accuracy.

#### G.3 Training instability

Our primary training run was performed before we introduced the changes described in Appendix B.2. While we have no reason to believe that the instabilities we observed there are a result of a bug in the OpenFold codebase, as opposed to an inherent limitation of the AlphaFold2 architecture, the former remains a possibility. It is unclear how potential issues of this kind may have affected runs that—like our primary training run—appeared to converge at the expected rate. [71] [72] [73] [74] [75]

## Footnotes

1 PDB accession code 7CJS_B ([38])

2 PDB accession code 7RRM_C ([39])

3 Predicted structure animations for diverse validation proteins can be accessed [here]. Residues are color-coded according to the 3-state secondary structure of the accompanying experimental fold.

4 For a comparison of the two techniques, see ([22]).

## References

[1] C. B. Anfinsen. Principles that govern the folding of protein chains. Science 181.4096 (1973), 223–230. 10.1126/science.181.4096.223.

[2] K. A. Dill, S. B. Ozkan, M. S. Shell, and T. R. Weikl. The protein folding problem. Annual Review of Biophysics 37 (2008), 289–316. 10.1146/annurev.bio-phys.37.092707.153558.

[3] D. T. Jones, T. Singh, T. Kosciolek, and S. Tetchner. MetaPSICOV: combining coevolution methods for accurate prediction of contacts and long range hydrogen bonding in proteins. Bioinformatics 31 (7 2015), 999–1006. 10.1093/bioinformatics/btu791.

[4] V. Golkov, M. J. Skwark, A. Golkov, A. Dosovitskiy, T. Brox, J. Meiler, and D. Cremers. Protein contact prediction from amino acid co-evolution using convolutional networks for graph-valued images. In: Advances in Neural Information Processing Systems. Ed. by D. Lee, M. Sugiyama, U. Luxburg, I. Guyon, and R. Garnett. Vol. 29. 2016. https://proceedings.neurips.cc/paper/2016/file/2cad8fa47bbef282badbb8de5374b894-Paper.pdf.

[5] S. Wang, S. Sun, Z. Li, R. Zhang, and J. Xu. Accurate De Novo Prediction of Protein Contact Map by Ultra-Deep Learning Model. PLOS Computational Biology 13.1 (2017), 1–34. 10.1371/journal.pcbi.1005324.

[6] Y. Liu, P. Palmedo, Q. Ye, B. Berger, and J. Peng. Enhancing Evolutionary Couplings with Deep Convolutional Neural Networks. Cell Systems 6 (1 2018), 65–74. 10.1016/j.cels.2017.11.014.

[7] A. W. Senior et al. Improved protein structure prediction using potentials from deep learning. Nature 577 (7792 2020), 706–710. 10.1038/s41586-019-1923-7.

[8] J. Xu, M. McPartlon, and J. Li. Improved protein structure prediction by deep learning irrespective of co-evolution information. Nature Machine Intelligence 3 (7 2021), 601–609. 10.1038/s42256-021-00348-5.

[9] A. Šali and T. L. Blundell. Comparative protein modelling by satisfaction of spatial restraints. Journal of Molecular Biology 234 (3 1993), 779–815. 10.1006/jmbi.1993.1626.

[10] A. Roy, A. Kucukural, and Y. Zhang. I-TASSER: a unified platform for automated protein structure and function prediction. Nature Protocols 5 (4 2010), 725–738. 10.1038/nprot.2010.5.

[11] J. Jumper et al. Highly accurate protein structure prediction with AlphaFold. Nature 577 (7792 2021), 583–589. 10.1038/s41586-021-03819-2.

[12] M. Mirdita, K. Schütze, Y. Moriwaki, L. Heo, S. Ovchinnikov, and M. Steinegger. ColabFold: making protein folding accessible to all. Nature Methods 19 (6 2022), 679–682. 10.1038/s41592-022-01488-1.

[13] M. Baek. Twitter post: Adding a big enough number for “residue_index” feature is enough to model hetero-complex using AlphaFold (green&cyan: crystal structure / magenta: predicted model w/ residue_index modification). 2021. https://twitter.com/minkbaek/status/1417538291709071362.

[14] T. Tsaban, J. K. Varga, O. Avraham, Z. Ben-Aharon, A. Khramushin, and O. SchuelerFurman. Harnessing protein folding neural networks for peptide–protein docking. Nature Communications 13 (1 2022), 176. 10.1038/s41467-021-27838-9.

[15] J. P. Roney and S. Ovchinnikov. State-of-the-art estimation of protein model accuracy using AlphaFold. bioRxiv (2022). 10.1101/2022.03.11.484043.

[16] A. Baltzis, L. Mansouri, S. Jin, B. E. Langer, I. Erb, and C. Notredame. Highly significant improvement of protein sequence alignments with AlphaFold2. Bioinformatics (2022). 10.1093/bioinformatics/btac625.

[17] P. Bryant, G. Pozzati, and A. Elofsson. Improved prediction of protein-protein interactions using AlphaFold2. Nature Communications 13 (1 2022), 1265. 10.1038/s41467022-28865-w.

[18] H. K. Wayment-Steele, S. Ovchinnikov, L. Colwell, and D. Kern. Prediction of multiple conformational states by combining sequence clustering with AlphaFold2. bioRxiv (2022). 10.1101/2022.10.17.512570.

[19] K. Tunyasuvunakool et al. Highly accurate protein structure prediction for the human proteome. Nature 596 (7873 2021), 590–596. 10.1038/s41586-021-03828-1.

[20] M. Varadi et al. AlphaFold Protein Structure Database: massively expanding the structural coverage of protein-sequence space with high-accuracy models. Nucleic Acids Research 50.D1 (2021), D439–D444. 10.1093/nar/gkab1061.

[21] E. Callaway. ‘The entire protein universe’: AI predicts shape of nearly every known protein. Nature 608 (4 2022), 15–16. 10.1038/d41586-022-02083-2.

[22] R. Evans, et al. Protein complex prediction with AlphaFold-Multimer. bioRxiv (2022). 10.1101/2021.10.04.463034.

[23] G. Ahdritz, N. Bouatta, S. Kadyan, L. Jarosch, D. Berenberg, I. Fisk, A. M. Watkins, S. Ra, R. Bonneau, and M. AlQuraishi. OpenProteinSet: Training data for structural biology at scale. 2023.

[24] A. Paszke, et al. PyTorch: An Imperative Style, High-Performance Deep Learning Library. In: Proceedings of the 33rd International Conference on Neural Information Processing Systems. 2019, 8026–8037. 10.5555/3454287.3455008.

[25] J. Bradbury et al. JAX: composable transformations of Python+NumPy programs. Version 0.3.13. 2018. http://github.com/google/jax.

[26] J. Rasley, S. Rajbhandari, O. Ruwase, and Y. He. DeepSpeed: System Optimizations Enable Training Deep Learning Models with Over 100 Billion Parameters. In: Proceedings of the 26th ACM SIGKDD International Conference on Knowledge Discovery & Data Mining. KDD ’20. 2020, 3505–3506. 10.1145/3394486.3406703.

[27] B. Charlier, J. Feydy, J. A. Glaunès, F.-D. Collin, and G. Durif. Kernel Operations on the GPU, with Autodiff, without Memory Overflows. Journal of Machine Learning Research 22.74 (2021), 1–6. http://jmlr.org/papers/v22/20-275.html.

[28] W. Falcon et al. PyTorch Lightning. Version 1.4. 2019. 10.5281/zenodo.3828935.

[29] T. Dao, D. Y. Fu, S. Ermon, A. Rudra, and C. Ré. FlashAttention: Fast and MemoryEfficient Exact Attention with IO-Awareness. 2022. 10.48550/ARXIV.2205.14135.

[30] M. Mirdita, L. von den Driesch, C. Galiez, M. J. Martin, J. Söding, and M. Steinegger. Uniclust databases of clustered and deeply annotated protein sequences and alignments. Nucleic Acids Research 45.D1 (2017), D170–D176. 10.1093/nar/gkw1081.

[31] wwPDB Consortium. Protein Data Bank: the single global archive for 3D macro-molecular structure data. Nucleic Acids Research 47.D1 (2018), D520–D528. 10.1093/nar/gky949.

[32] J. Haas, A. Barbato, D. Behringer, G. Studer, S. Roth, M. Bertoni, K. Mostaguir, R. Gumienny, and T. Schwede. Continuous Automated Model EvaluatiOn (CAMEO) complementing the critical assessment of structure prediction in CASP12. Proteins: Structure, Function, and Bioinformatics 86 (Suppl 1 2018), 387–398. 10.1002/prot.25431.

[33] V. Mariani, M. Biasini, A. Barbato, and T. Schwede. lDDT: a local superposition-free score for comparing protein structures and models using distance difference tests. Bioinformatics 29 (21 2013), 2722–2728. 10.1093/bioinformatics/btt473.

[34] H. L. Knox, E. K. Sinner, C. A. Townsend, A. K. Boal, and S. J. Booker. Structure of a _1_ -dependent radical SAM enzyme in carbapenem biosynthesis. Nature 602 (7896 2022), 343–348. 10.1038/s41586-021-04392-4.

[35] C. A. Orengo, A. D. Michie, S. Jones, D. T. Jones, M. B. Swindells, and J. M. Thornton. CATH–a hierarchic classification of protein domain structures. Structure 5 (8 1997), 1093–1108. 10.1016/s0969-2126(97)00260-8.

[36] I. Sillitoe et al. CATH: increased structural coverage of functional space. Nucleic Acids Research 49.D1 (2021), D266–D273. 10.1093/nar/gkaa1079.

[37] A. Andreeva, E. Kulesha, J. Gough, and A. G. Murzin. The SCOP database in 2020: expanded classification of representative family and superfamily domains of known protein structures. Nucleic Acids Research 48.D1 (2020), D376–D382. 10.1093/nar/gkz1064.

[38] Y. Saitoh et al. Structural basis for high selectivity of a rice silicon channel Lsi1. Nature Communications 12 (1 2021), 6236. 10.1038/s41467-021-26535-x.

[39] D. C. A. M. Mota, I. A. Cardoso, R. M. Mori, M. R. B. Batista, L. G. M. Basso, C. M. Nonato, A. J. Costa-Filho, and L. F. S. Mendes. Structural and thermodynamic analyses of human TMED1 (p24 1) Golgi dynamics. Biochimie 192 (2022), 72–82. 10.1016/j.biochi.2021.10.002.

[40] P. Koehl. Protein structure similarities. Current Opinion in Structural Biology 11 (3 2001), 348–353. 10.1016/s0959-440x(00)00214-1.

[41] M. AlQuraishi. End-to-End Differentiable Learning of Protein Structure. Cell Systems 8.4 (2019), 292–301.e3. 10.1016/j.cels.2019.03.006.

[42] W. Kabsch and C. Sander. Dictionary of protein secondary structure: pattern recognition of hydrogen-bonded and geometrical features. Science 22 (12 1983), 2577–2637. 10.1002/bip.360221211.

[43] A. Zemla. LGA: a method for finding 3D similarities in protein structures. Nucleic Acids Research 31 (13 2003), 3370–3374. 10.1093/nar/gkg571.

[44] A. Vaswani, N. Shazeer, N. Parmar, J. Uszkoreit, L. Jones, A. N. Gomez, L. Kaiser, and I. Polosukhin. Attention Is All You Need. 2017. 10.48550/ARXIV.1706.03762.

[45] M. N. Rabe and C. Staats. Self-attention Does Not Need *O*(*n*^2^) Memory. 2021. 10.48550/ARXIV.2112.05682.

[46] S. Cheng, R. Wu, Z. Yu, B. Li, X. Zhang, J. Peng, and Y. You. FastFold: Reducing Al-phaFold Training Time from 11 Days to 67 Hours. 2022. 10.48550/ARXIV.2203.00854.

[47] Z. Li, X. Liu, W. Chen, F. Shen, H. Bi, G. Ke, and L. Zhang. Uni-Fold: An Open-Source Platform for Developing Protein Folding Models beyond AlphaFold. bioRxiv (2022). 10.1101/2022.08.04.502811.

[48] D. S. Marks, L. J. Colwell, R. Sheridan, T. A. Hopf, A. Pagnani, R. Zecchina, and C. Sander. Protein 3D Structure Computed from Evolutionary Sequence Variation. PLOS ONE 6.12 (2011), 1–20. 10.1371/journal.pone.0028766.

[49] J. I. Sułkowska, F. Morcos, M. Weigt, T. Hwa, and J. Onuchic. Genomics-aided structure prediction. Proceedings of the National Academy of Sciences 109 (26 2012), 10340–10345. 10.1073/pnas.1207864109.

[50] J. Kaplan, S. McCandlish, T. Henighan, T. B. Brown, B. Chess, R. Child, S. Gray, A. Radford, J. Wu, and D. Amodei. Scaling Laws for Neural Language Models (2020). 10.48550/ARXIV.2001.08361.

[51] J. Hoffmann et al. Training Compute-Optimal Large Language Models (2022). 10.48550/ARXIV.2203.15556.

[52] Y. Tay, M. Dehghani, S. Abnar, H. W. Chung, W. Fedus, J. Rao, S. Narang, V. Q. Tran, D. Yogatama, and D. Metzler. Scaling Laws vs Model Architectures: How does Inductive Bias Influence Scaling? (2022). 10.48550/ARXIV.2207.10551.

[53] G. E. Karniadakis, I. G. Kevrekidis, L. Lu, P. Perdikaris, S. Wang, and L. Yang. Physics-informed machine learning. Nature Reviews Physics 3 (6 2021), 422–440. 10.1038/s42254-021-00314-5.

[54] Z. Lin et al. Evolutionary-scale prediction of atomic-level protein structure with a language model. Science 379.6637 (2023), 1123–1130. 10.1126/science.ade2574.

[55] E. C. Alley, G. Khimulya, S. Biswas, M. AlQuraishi, and G. M. Church. Unified rational protein engineering with sequence-based deep representation learning. Nature Methods 16 (12 2019), 1315–1322. 10.1038/s41592-019-0598-1.

[56] R. Chowdhury et al. Single-sequence protein structure prediction using a language model and deep learning. Nature Biotechnology (2022). 10.1038/s41587-022-01432-w.

[57] R. Wu, et al. High-resolution de novo structure prediction from primary sequence. bioRxiv (2022). 10.1101/2022.07.21.500999.

[58] J. Singh, K. Paliwal, T. Litfin, J. Singh, and Y. Zhou. Predicting RNA distance-based contact maps by integrated deep learning on physics-inferred secondary structure and evolutionary-derived mutational coupling. Bioinformatics 38.16 (2022), 3900–3910. 10.1093/bioinformatics/btac421.

[59] M. Baek, R. McHugh, I. Anishchenko, D. Baker, and F. DiMaio. Accurate prediction of nucleic acid and protein-nucleic acid complexes using RoseTTAFoldNA. bioRxiv (2022).

[60] R. Pearce, G. S. Omenn, and Y. Zhang. De Novo RNA Tertiary Structure Prediction at Atomic Resolution Using Geometric Potentials from Deep Learning. bioRxiv (2022). 10.1101/2022.05.15.491755.

[61] M. McPartlon, B. Lai, and J. Xu. A Deep SE(3)-Equivariant Model for Learning Inverse Protein Folding. bioRxiv (2022). 10.1101/2022.04.15.488492.

[62] M. McPartlon and J. Xu. An end-to-end deep learning method for rotamer-free protein side-chain packing. bioRxiv (2022). 10.1101/2022.03.11.483812.

[63] Y. Zhang and J. Skolnick. Scoring function for automated assessment of protein structure template quality. Proteins 57 (4 2004), 702–710. 10.1002/prot.20264.

[64] S. Rajbhandari, J. Rasley, O. Ruwase, and Y. He. ZeRO: Memory Optimizations To-ward Training Trillion Parameter Models. 2019. 10.48550/ARXIV.1910.02054.

[65] D. P. Kingma and J. Ba. Adam: A Method for Stochastic Optimization. In: 3rd International Conference on Learning Representations, ICLR 2015, San Diego, CA, USA, May 7-9, 2015, Conference Track Proceedings. 2015. http://arxiv.org/abs/1412.6980.

[66] G. Wang, X. Fang, Z. Wu, Y. Liu, Y. Xue, Y. Xiang, D. Yu, F. Wang, and Y. Ma. HelixFold: An Efficient Implementation of AlphaFold2 using PaddlePaddle. 2022. 10.48550/ARXIV.2207.05477.

[67] J. Yuan, et al. OneFlow: Redesign the Distributed Deep Learning Framework from Scratch. 2021. 10.48550/ARXIV.2110.15032.

[68] K. Hinsen, S. Hu, G. R. Kneller, and A. J. Niemi. A comparison of reduced coordinate sets for describing protein structure. Journal of Chemical Physics 139 (12 2013), 124115. 10.1063/1.4821598.

[69] A. Krokhotin, A. Liwo, A. J. Niemi, and H. A. Scheraga. Coexistence of Phases in a Protein Heterodimer. Journal of Chemical Physics 137.3 (2012), 035101. 10.1063/1.4734019.

[70] S. Ovchinnikov. Twitter post: Weekend project! So now that OpenFold weights are available. I was curious how different they are from AlphaFold weights and if they can be used for AfDesign evaluation. More specifically, if you design a protein with AlphaFold, can OpenFold predict it (and vice-versa)? (1/5). 2022. https://twitter.com/sokrypton/status/1551242121528520704.

[71] T. Dang et al. Molecular basis of antibiotic self-resistance in a bee larvae pathogen. Nature Communications 13 (1 2022), 2349. 10.1038/s41467-022-29829-w.

[72] B. Huang, Y. Xu, X. Hu, Y. Liu, S. Liao, J. Zhang, C. Huang, J. Hong, Q. Chen, and H. Liu. A backbone-centred energy function of neural networks for protein design. Nature 602 (7897 2022), 523–528. 10.1038/s41586-021-04383-5.

[73] X. Wei et al. The α-Helical Cap Domain of a Novel Esterase from Gut Alistipes shahii Shaping the Substrate-Binding Pocket. Journal of Agricultural Food chemistry 69 (21 2021), 6064–6072. 10.1021/acs.jafc.1c00940.

[74] A. C. Y. Yu, G. Volkers, S. A. K. Jongkees, L. J. Worrall, S. G. Withers, and N. C. J. Strynadka. Crystal structure of the Propionibacterium acnes surface sialidase, a drug target for P. acnes-associated diseases. Glycobiology 32.2 (2021), 162–170. 10.1093/gly-cob/cwab094.

[75] B. L. Carroll, K. E. Zahn, J. P. Hanley, S. S. Wallace, J. A. Dragon, and S. Doublié. Caught in motion: human NTHL1 undergoes interdomain rearrangement necessary for catalysis. Nucleic Acids Research 49 (22 2021), 13165–13178. 10.1093/nar/gkab1162.

